# Dynamic translocation of Inside-Out proteins to the cell surface underlies cellular adaptation to cancer-induced stress

**DOI:** 10.1101/2025.10.30.685532

**Authors:** Tomasz Slezak, Kelly M. O’Leary, Tanya Guevara Avella, Natalia Musial, Jinyang Li, Anna Andrzejczak, Elizabeth F. Scott, Duc Anh Le, Anthony A. Kossiakoff

**Affiliations:** Department of Biochemistry and Molecular Biology, The University of Chicago, Chicago, IL 60637; Intercollegiate Faculty of Biotechnology, University of Gdansk and Medical University of Gdansk, Poland; Department of Experimental Therapy, Hirszfeld Institute of Immunology and Experimental Therapy, Polish Academy of Sciences, Poland; Institute for Biophysical Dynamics, The University of Chicago, Chicago, IL 60637

## Abstract

Inside-out (I-O) protein display, the non-canonical surface localization of intracellular proteins, represents an underexplored feature of tumor cell biology. Here, we map the molecular landscape and trafficking mechanisms that control the presentation of I-O proteins on cancer cell membranes. Employing APEX2-mediated proximity biotinylation and a custom antibody generation and validation platform, we identified approximately 140 high-confidence I-O proteins, primarily ribosomal, proteasomal, chaperone, and translation factors, notably enriched in protein families associated with stress-response pathways. Validation of 500 antibodies encompassing 40 I-O targets across seven tumor cell lines confirmed selective and robust surface localization, while *in vivo* imaging in mouse xenografts demonstrated pronounced and tumor-specific antibody accumulation. I-O proteins were absent on PBMCs and in normal tissues, indicating cancer cell selectivity. Functional analyses revealed that I-O protein tethering to the membrane is dependent on heparan sulfate interactions; enzymatic removal of these glycans led to the clearance of I-O proteins from the cell surface. Notably, the removed proteins returned to baseline levels within six hours, indicating a dynamic balance related to ER-Golgi trafficking and cellular stress. Nearly half of these I-O proteins overlapped with known stress granule components; however, stress elements that promote stress granule formation do not similarly affect surface display of I-O proteins. Furthermore, I-O proteins are present on standard cancer cell lines under lower stress levels needed to induce stress granule formation, suggesting parallel yet mechanistically distinct aspects of the stress response. These findings position I-O display as a new paradigm in protein trafficking, different from traditional secretion pathways and closely linked to stress response.

## Introduction

The plasma membrane serves as a selectively permeable barrier, mediating the transduction of extracellular signals into specific intracellular responses and thereby playing a fundamental role in cellular signaling. Its structural architecture ensures selective trafficking of molecules, maintaining the integrity of the intracellular environment by permitting only suitable substances to cross while preventing the entry of harmful agents or loss of essential cellular components. While most external stimuli are detected and relayed via transmembrane receptors or transporters, many proteins of extracellular origin are internalized through well-characterized endocytic pathways to fulfill essential intracellular functions (1). Conversely, numerous intracellular proteins can be exported and released into the extracellular space via vesicles or exosomes, where they participate in intercellular communication (2).

Unlike these mechanisms, there exists a less well-characterized subset of proteins, herein referred to as “inside-out” (I-O) proteins. These proteins are primarily associated with intracellular functions; however, under specific cellular conditions, they undergo translocation and are presented on the external surface of the plasma membrane (3–6). A notable characteristic of I-O proteins is the absence of canonical secretory signals or other membrane-targeting motifs that would typically facilitate their localization to the external leaflet of the plasma membrane. Consequently, the identification of novel members demonstrating inside-out behavior has likely been hindered by their inconsistency with the prevailing dogma concerning protein translocation mechanisms. As a result, the unforeseen presence of these potential I-O proteins may have previously been disregarded as experimental artifacts (7). Nevertheless, as additional instances have been documented and validated through increasingly rigorous scientific methodologies, the recognition of this class of proteins has become widely accepted.

Notably, several of the earliest identified I-O proteins exhibit dual, context-dependent functions, referred to as “moonlighting” activities, differing between their intracellular and membrane-associated states. Heat shock proteins, particularly members of the Hsp70 family, exemplify this phenomenon (6, 8–12); beyond their well-known intracellular chaperone roles, surface-localized Hsp70 proteins stabilize and protect stressed plasma membranes, function as immunological danger signals, modulate cell signaling pathways, and facilitate intercellular communication via vesicular export (13). Another prominent example is extracellular nucleolin, which acts as a multifunctional cell surface receptor for nucleic acids and other ligands, thereby influencing immune activation and the regulation of inflammation (4, 14). Additional I-O proteins with established intracellular and membrane-associated roles include metabolic enzymes, elongation factors, and cytoskeletal proteins (15– 17). Recent advances in cell surface proteomics have highlighted the unexpected presence of RNA-associated intracellular proteins displayed on the plasma membrane (18–20). Yet, the scope, regulation, and functional implications of this phenomenon in tumor biology are only beginning to be explored.

To more broadly map the I-O protein landscape, we employed an APEX2-mediated proximity biotinylation approach in concert with targeted antibody screening. This integrated strategy enabled us not only to identify but also to validate the occurrence, kinetics, and selectivity of a cohort of approximately 140 surface-displayed I-O proteins in tumor cells, a set spanning diverse functional classes such as ribosomal and proteasomal subunits, heat shock proteins, and translation machinery. Gene ontology enrichment revealed a pronounced association with stress-adaptive pathways, implicating surface presentation of these proteins as a key facet of cellular stress responses.

To substantiate these findings and enhance their biological relevance, we developed a high-throughput panel of nearly 500 antibodies targeting 40 representative I-O antigens across various functional phenotypes. Multiplexed antibody profiling across seven cancer cell lines revealed celltype-dependent abundance of these proteins at the cell surface, findings further confirmed by *in vivo* imaging in a breast cancer xenograft model. Notably, the surface accumulation of I-O proteins was highly tumor-specific; normal tissue and unstressed immune cells did not exhibit detectable surface display. Conversely, exposure of peripheral blood mononuclear cells (PBMCs) to pharmacological stress significantly induced I-O presentation, an effect that was reversible upon alleviation of stress. These findings highlight the dynamic and stress-responsive nature of I-O protein trafficking to the plasma membrane.

The fact that I-O proteins are synthesized as soluble forms without prominent membrane attachment motifs raises questions about how they are tightly retained on the membrane surface. We show that proteinase K digestion selectively removes these proteins from the cell surface (21, 22). Further enzymatic and chemical tests identified heparan sulfate glycosaminoglycans as a key mediator of I-O membrane tethering, with Heparinase II treatment effectively removing virtually all surveyed I-O proteins from the cells. Notably, surface display was quickly restored after enzymatic removal, as long as ER-Golgi trafficking remained functional and cellular processes were not inhibited by low temperature or pharmacological agents. This indicates a highly dynamic, trafficking-dependent balance between the surface and internal pools of I-O proteins. The underlying mechanisms that regulate the levels of these pools to maintain this homeostasis still need to be determined.

Notably, the families and functional categories present within the I-O proteome closely resemble those characterizing cytoplasmic stress granules (SGs), which are pivotal to cellular adaptation and fate determination under adverse conditions (23). Comparative analyses have determined that nearly fifty percent of the surface I-O proteins coincide with recognized SG components; however, this figure likely underestimates the actual overlap, owing to disparities in proteome cataloging. Notwithstanding these similarities, I-O proteins are observed on the surface as dispersed, folded entities tethered by glycosaminoglycans, rather than the internally organized assemblies that are characteristic of SGs (23, 24). Another key distinction is that I-O proteins act as early responders to cellular stress, consistently present at baseline on the surfaces of standard cancer cell lines. In contrast, SGs are typically absent under basal conditions in cancer cell lines and only form when cells are subjected to heightened stress (25, 26).

Together, these findings reveal that surface-presented I-O proteins and stress granule components constitute distinct yet interconnected arms of the stress response, the former functioning as a rapidly mobilized, membrane-accessible pool dynamically regulated by stress stimuli and trafficking pathways. The absence of canonical secretion or membrane-localization signals among I-O proteins raises intriguing questions about non-canonical trafficking and the broader implications of this pathway for tumor biology, cellular homeostasis, and immune surveillance. Understanding how cells coordinate these dual adaptive programs will likely open new opportunities for exploring stress biology and targeting tumor-specific vulnerabilities in cancer therapy.

## Results

### Identification of I-O proteins on the cell surface

Selective identification of I-O proteins on the surface of HCT116 cells was accomplished using APEX2-mediated labeling combined with mass spectrometry. APEX2, an engineered ascorbate peroxidase, produces short-lived biotinphenoxyl radicals upon addition of biotin-PEG4-phenol and H_2_O_2_ to facilitate the selective labeling of proximal tyrosine residues (27–29). Importantly, these radicals are membrane-impermeable, ensuring exclusive labeling of extracellular proteins and avoiding contamination from intracellular sources (Fig. S1) (30).

In an effort to enhance the signal-to-noise ratio, we conducted a comparative analysis between untreated HCT116 cells and those subjected to mild treatment with proteinase K (21, 22, 31). This procedure selectively removed the majority of I-O proteins without significantly affecting the overall membrane proteome (Fig. 1A). The resulting “shaved” dataset facilitated unambiguous identification of likely I-O proteins, as illustrated in the volcano plot in Fig. 1B. Nonetheless, certain limitations may give rise to false negatives. The presentation of surface-expressed individual I-O proteins may vary depending on the cell line, which could result in the failure to identify certain I-O proteins of lower abundance, contingent upon the cell line employed in the experiment, in this case, HCT116. Labeling efficiency tends to favor larger proteins containing multiple exposed tyrosines. Proteins lacking tyrosines or possessing tyrosines concealed by tertiary structure, protein interactions, or membrane associations may evade detection under strict cutoff criteria. Additionally, we observed a few instances where enzymatic treatment inadvertently removed potential biotinylation sites on cognate membrane proteins, leading to false positives; however, these were readily curated and excluded from the dataset. Although relaxing these thresholds increased the number of identified I-O proteins, our primary objective was to define general families and functions rather than to exhaustively catalog the entire I-O proteome. Based on these criteria, approximately 140 I-O candidates were identified falling into a number of distinct protein function families related to the stress response (Fig. 1C, 1D).

**Figure 1.**
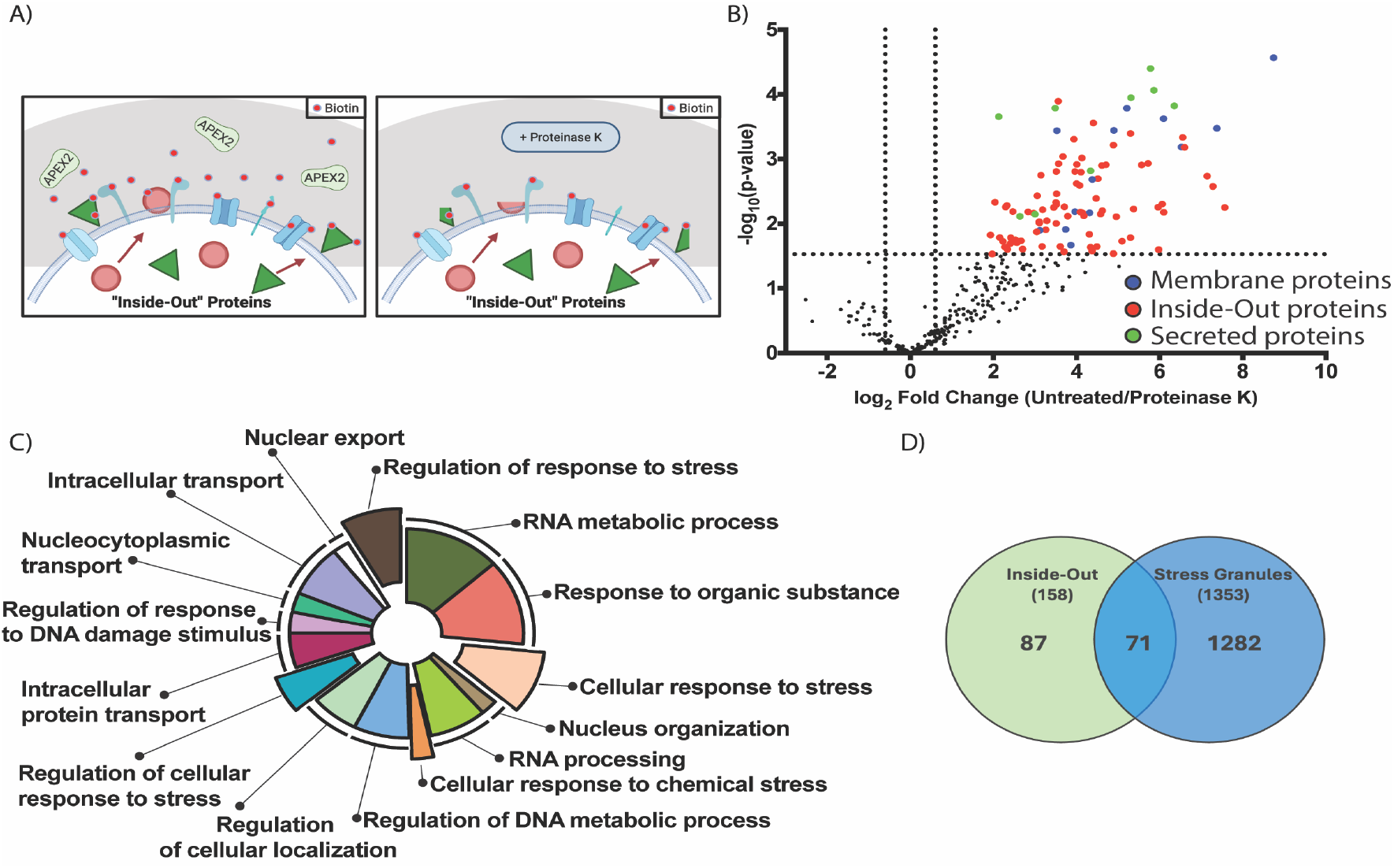
The discovery of Inside-Out proteins. **(A)** Schematic illustrating APEX2-mediated labeling of Inside-Out proteins on the cell surface. Utilizing APEX2 and a non-permeabilizing biotin substrate, the cell surface was efficiently labeled and subsequently digested using proteinase K. **(B)** Volcano plot of enriched proteins resulting from a comparative APEX2 mass spectrometry analysis between untreated and proteinase K-treated cells. Cell surface proteins exhibiting the greatest sensitivity to proteinase K digestion are located in the upper right quadrant (n = 3 biological replicates). **(C)** Enrichment analysis visualization of the biological processes (based on Gene Ontology) for statistically significant proteins. A large subset of identified proteins is responsible for the cellular stress response. **(D)** Venn diagram illustrating the overlap between identified Inside-Out proteins and proteins commonly found in canonical intracellular stress granules. Nearly half of the identified Inside-Out proteins are also present in stress granules.

Target-specific antibodies are regarded as the gold standard for profiling cell surface proteins and were employed herein to validate the surface display of candidate I-O targets. To generate synthetic antibodies targeting selected I-O proteins, a high-throughput phage display mutagenesis pipeline, previously described (32, 33), was utilized. These antibodies are derived from Fab phage display synthetic libraries employing a reduced genetic code strategy (34). An initial selection of 67 candidate I-O proteins was conducted for antibody generation, based on multiple criteria: top hits from proteomics data, mid-range candidates, and lowerabundance proteins from families with known I-O relevance. Antigen design also considered the feasibility of producing high-quality, representative protein domains, particularly for large or intrinsically disordered proteins. In such instances, stable regions were identified based on structural data or AlphaFold predictions. While some antigens were produced in *E. coli*, others necessitated mammalian expression and refolding procedures. Due to issues related to expression or stability, 17 antigens were excluded, resulting in 50 targets deemed suitable for phage display selection.

### Phage Display Selection and Characterization of anti-I-O Antigen Fabs

The 50 I-O antigens underwent four to five iterative rounds of phage display selection, with progressively decreasing antigen concentrations to augment selection stringency. In the final round, antigens were typically employed at a concentration of 10 nM, resulting in enriched pools of high-affinity binders, as confirmed by phage ELISA. Unique clones were identified through sequencing, and approximately 500 antibody fragments (Fabs) were cloned, expressed, and purified for subsequent validation. We note that the Fabs produced by these procedures recognize fully folded forms of the antigens they are derived from, whether in their soluble or membrane-associated states.

Initial screening utilizing single-point protein ELISAs against the specific soluble antigens demonstrated that the majority of Fabs exhibited binding affinities within the low-to mid-nanomolar range (Fig. S2). However, ten I-O targets yielded few or no Fabs meeting this criterion and were consequently excluded. The remaining 40 targets were subsequently assessed for surface binding via flow cytometry with non-permeabilized HCT116 cells. Representative binding histograms are presented in Fig. 2A, S3. Notably, there was a weak correlation between an antibody’s binding to soluble antigen and its ability to recognize the same antigen in a cell surface context. For many of the targets, fewer than half of the Fabs showed detectable binding on the cell surface, and those that did often exhibited reduced binding compared to their interaction with soluble antigen. These results suggest that spatial factors, such as epitope orientation or membrane-dependent structural features, may substantially influence surface recognition. Furthermore, cell surface immunofluorescence staining of HCT116 cells with a panel of Fab fragments targeting the 40 I-O targets showed variably dispersed surface localization along the external membranes, without evidence of clear concentration into large, condensed structures (Fig. 2B, S4). However, the limited resolution of these images does not rule out this possibility.

**Figure 2.**
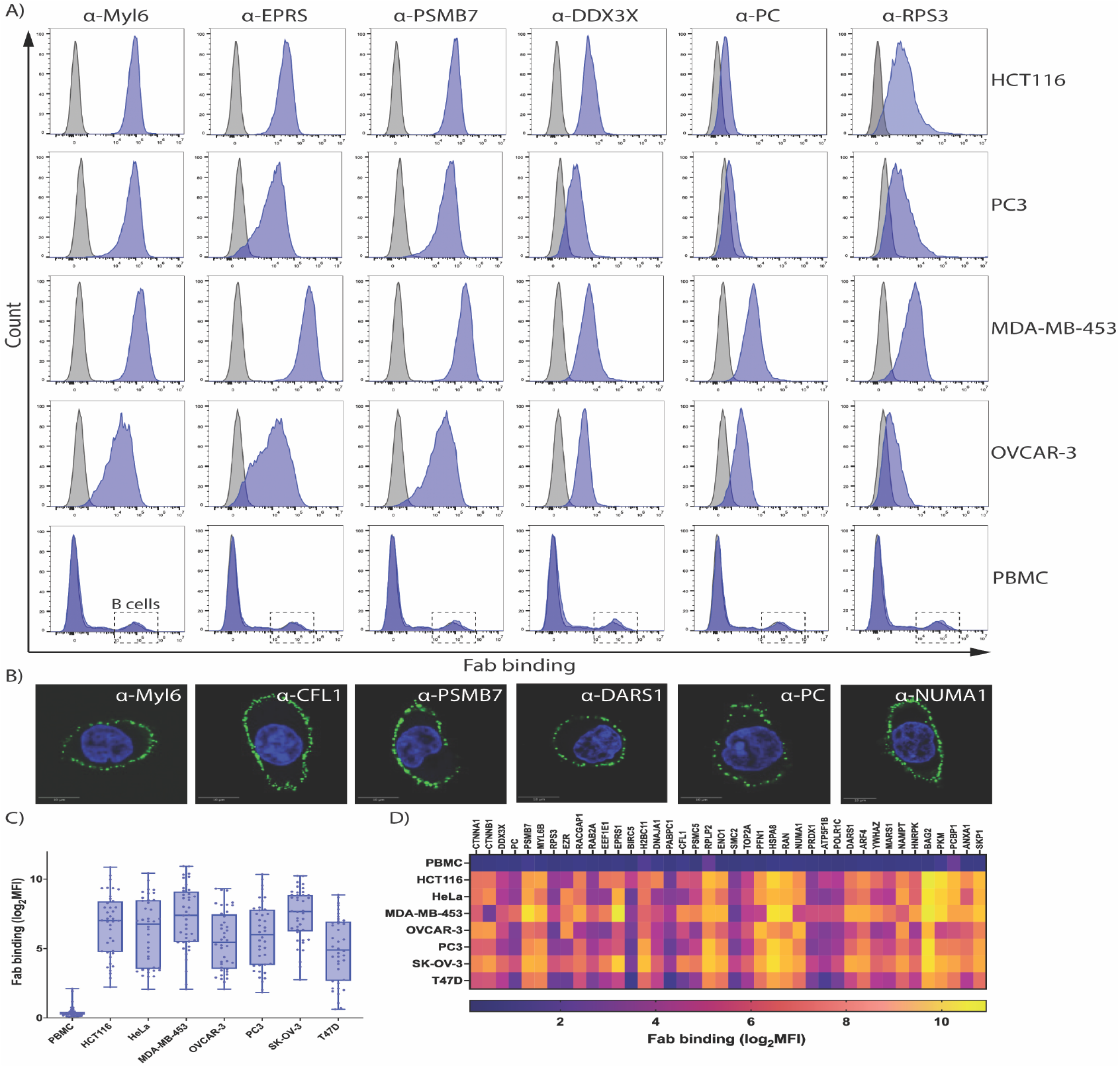
Cell surface detection of Inside-Out proteins using antibody-based staining. **(A)** Representative flow cytometry profiles demonstrating surface staining of various cancer cell lines and healthy PBMCs with antibodies against Inside-Out proteins (purple) compared to secondary antibody only (gray). **(B)** Representative confocal microscopy images showing HCT116 live cell surface staining with antibodies targeting InsideOut proteins (green) and nuclei staining with DAPI (blue). **(C)** Box plot quantifying antibody binding to the cell surface across multiple cancer cell lines for 40 Inside-Out proteins. No surface binding is observed in healthy PBMCs. Scale bar denotes 10 *μ*m. **(D)** Heat map depicting the surface expression of 40 Inside-Out proteins across eight distinct cell types. Each column represents an individual Inside-Out protein, and each row corresponds to a specific cell type. The color scale reflects the extent of antibody-mediated surface binding.

Based on the performance of flow cytometry in HCT116 cells and the EC_50_ binding data, the 2-3 top-performing Fabs for each I-O target were selected for extended analysis across six additional cancer cell lines. To reduce experimental variability, all Fabs were tested simultaneously for each cell type within a single flow cytometry run. Fig. 2C presents box plots illustrating the distributions of mean fluorescence intensity (MFI) for the group of I-O targets across the panel of seven cancer cell lines. While some comparisons of MFI between targets may be affected by differences in Fab affinity and epitope accessibility, the relative levels provide valuable insights into target-specific variations in surface display abundance. This is further highlighted by the heat map (Fig. 2D), which reveals overall patterns of relative surface expression across all 40 I-O targets in seven cell lines, showing significant variability in display levels for several targets.

The heterogeneity observed in surface staining patterns among various cell types underscores the importance of assessing multiple cell lines to determine the relative surface display levels of antigens. Fig. 2A illustrates representative flow cytometry histograms obtained from four cancer cell lines, which were analyzed with Fab fragments targeting six functionally diverse I-O markers: Myl6 (myosin light chain 6), EPRS (glutamyl-prolyl-tRNA synthetase), PSMB7 (proteasome subunit β7), DDX3X (DEAD-box helicase 3), PC (pyruvate carboxylase), and RPS3 (40S ribosomal protein S3). All six antigens were detectable on the cell surface, albeit with varying intensities. Myl6, EPRS, and PSMB7 exhibited strong staining across all four cell lines. DDX3X was identified in three lines, with minimal signal detected in PC3 cells. Conversely, PC and RPS3 displayed more restricted binding, observable in only two cell lines.

To evaluate whether the surface display of I-O antigens is a stress-related phenomenon associated with cancer cells, normal primary peripheral blood mononuclear cells (PBMCs) were employed as a surrogate for non-transformed cells. Flow cytometry analysis using the same panel of Fab binders demonstrated no detectable binding above background levels (Fig. 2A), indicating that the surface presentation of I-O antigens does not occur under basal conditions. This investigation was further expanded to include the entire set of I-O Fabs, none of which exhibited binding to PBMCs (Fig. 2D), thereby supporting the hypothesis that surface localization is stress-induced.

To further investigate whether stress-inducing stimuli in non-cancerous cells could replicate this effect, we treated PBMCs with GW4869, an inhibitor of neutral sphingomyelinase known to elicit cellular stress responses and inhibit the formation of exosomes (35). After one hour of treatment, PBMCs were analyzed by flow cytometry using a representative panel of Fab binders. Notably, all tested Fabs displayed measurable binding, suggesting that GW4869-induced stress was sufficient to drive surface localization of I-O antigens in otherwise normal cells (Fig. S5). Importantly, following a 24-hour recovery period without the inhibitor, Fab binding returned to baseline, and PBMCs reverted to their unstressed phenotype. These results indicate that surface display of I-O antigens is a reversible, stress-dependent event, and may serve as a marker of the cellular stress state characteristic of cancer or other diseases. Notably, treating PBMC cells with sodium arsenite and the mTOR inhibitor, Torin 1, (both known to induce stress granule formation in cancer cell lines (36, 37)) did not induce surface display of I-O proteins on PBMCs, unlike GW4869 (Fig. S6). This suggests that the display of I-O proteins on the cell surface is triggered by mechanisms different from those that lead to stress granule formation.

### Tumor-specific surface display of Inside-Out protein enables selective antibody accumulation in xenograft models

Building upon *in vitro* evidence demonstrating robust antibody binding to tumor cell surfaces, we investigated the *in vivo* surface display of I-O antigens on cancer and normal cells using a xenograft murine model. Numerous I-O antigens possess highly conserved sequences, reflecting the fundamental cellular functions preserved across eukaryotic species. We selected six I-O targets previously characterized by flow cytometry, chosen for their high sequence homology between human and mouse (>96% identity): Myl6 (100%), RPS3 (100%), EPRS (96%), PC (98%), DDX3X (99%), and PSMB7 (99%). This degree of conservation facilitated the assessment of tumor-targeting efficacy and off-target binding of I-O–specific antibodies *in vivo*.

To monitor antibody localization, Fab fragments targeting each I-O antigen were indirectly labeled with Cy7 using an engineered Protein G domain (PGA1-Cy7), which binds to the engineered Fab scaffold with high affinity in a modular, plug-and-play manner (38). HCT116 tumor xenografts were established subcutaneously in immunodeficient nude mice and allowed to grow for six days before intravenous administration of fluorescently labeled Fabs. *In vivo* fluorescence imaging conducted at 2and 3-hours post-injection demonstrated significant, tumor-specific antibody accumulation, which persisted for at least 190 minutes, with clear retention within the tumor microenvironment (Fig. 3A, B). To confirm antibody selectivity, we performed flow cytometry analyses on normal mouse kidney cells using anti-PC and anti-PSMB7 Fabs. No binding was detected, consistent with previous findings in PBMCs, indicating these IO antigens are not displayed on the surface of non-tumor cells (Fig. 3C). Overall, these data show the targeted *in vivo* delivery of conserved I-O proteins to tumors, supporting their potential as cancer-specific therapeutic targets.

**Figure 3.**
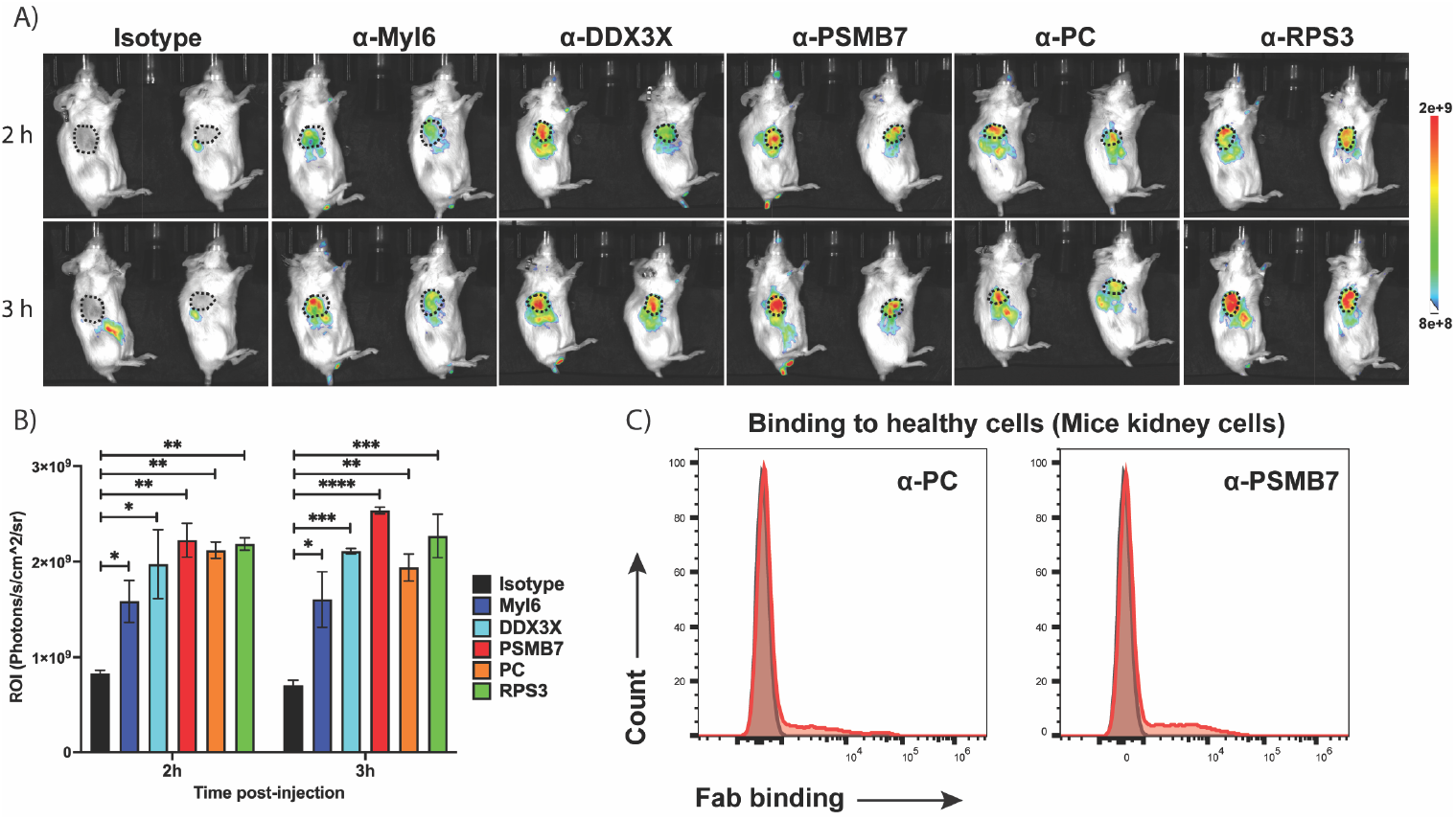
In vivo validation of tumor-specific accumulation of antibodies targeting Inside-Out proteins. **(A)** In vivo tumor accumulation of antibodies against Inside-Out proteins was assessed using a mouse model bearing HCT116 cancer xenografts. Fluorescently labeled antibodies were administered intravenously for the indicated time points. Robust and selective accumulation within the tumor microenvironment was observed as indicated by strong fluorescent signals localized to tumor sites (n = 2 mice). **(B)** Quantification of antibody accumulation within the tumor microenvironment from the experiment in A (n = 2, mean ± SD. *P ≤ 0.05, **P ≤ 0.01, ***P ≤ 0.001, ****P ≤ 0.0001. Two-way ANOVA) **(C)** Representative flow cytometry analysis showing the absence of binding by antibodies against PC and PSMB7 (red) to healthy cells isolated from mouse kidney, indicating minimal off-target interactions.

### Mechanism of Adhesion of I-O Protein to the Cell Surface

I-O proteins do not possess intrinsic membrane-targeting motifs or recognized post-translational modifications that could explain their prominent localization at the plasma membrane. Given their diverse biophysical properties, it was deemed important to determine how I-O proteins interact with heterogeneous membrane surfaces-either as individual entities or when associated with other molecules. Previous studies indicated that treatment with proteinase K effectively removes many I-O proteins from the cell surface (39); however, due to its broad substrate specificity, this effect may reflect the digestion of the I-O proteins themselves, associated matrix components, or both. To gain a clearer mechanistic understanding, we explored less invasive treatments to clarify how I-O proteins are anchored to the membrane.

We utilized the same panel of 40 Fab antibodies targeting I-O markers to assess changes in surface staining through flow cytometry after interventions aimed at disrupting specific classes of cell surface interactions in HCT116 cells (Fig. 4A). Treatment with 0.5 M NaCl, intended to interfere with electrostatic forces, resulted in only a slight reduction in surface Fab binding across the panel, suggesting that simple ionic interactions are unlikely to mediate I-O protein association. Similarly, treatment with PNGase F, which removes N-linked glycans, showed no significant effect. Given the enrichment of nucleic acid-binding proteins among I-O antigens and prior evidence of extracellular RNA on the surfaces of cancer cells (19, 20), we further examined the effect of RNase A digestion. Of the antigen panel tested, six I-O proteins were associated with RNA-related functions or binding. The surface display of these proteins was markedly decreased following enzymatic digestion (Fig. S7), whereas I-O proteins not involved in RNA functions did not show similar changes. This observation suggests that the latter group of proteins attaches to the membrane surface through alternative mechanisms.

**Figure 4.**
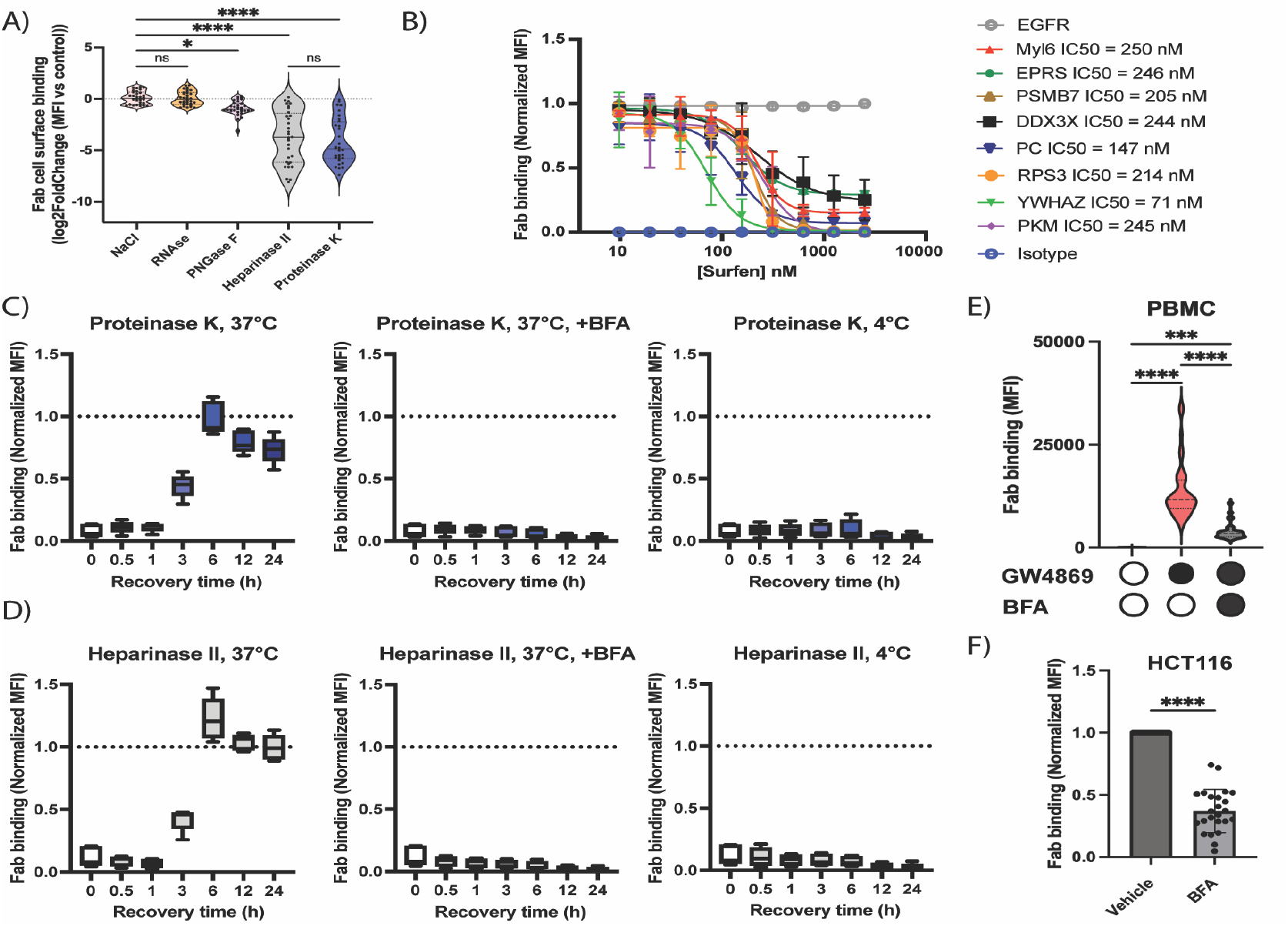
Selective removal and recovery of Inside-Out proteins from cancer cell surfaces following enzymatic and chemical treatments. (**A)** Violin plot of Inside-Out protein surface abundance on HCT116 cells after treatments with 0.5 M NaCl for 1 minute (n = 40), RNase A for 1 hour (n = 40), PNGase F for 1 hour (n = 40), Heparinase II for 15 minutes (n = 40), and Proteinase K for 1 hour (n = 40). Marked reductions were observed after Heparinase II and Proteinase K treatment, suggesting interactions between Inside-Out proteins and heparan sulfate. (n = 40, *\*P ≤* 0.05, \**\*P ≤* 0.01, \*\**\*P ≤* 0.001, \*\*\**\*P ≤* 0.0001. Two-way ANOVA). **(B)** Dose-dependent displacement of Inside-Out proteins by surfen. Increasing concentrations of surfen abolished Fab binding across all tested Inside-Out antibodies with similar IC_50_ values, while binding of an EGFR antibody remained unaffected. Fab binding was normalized to the untreated control (n = 3, mean ± SD). **(C)** Recovery of Inside-Out protein display after Proteinase K treatment. Following 1 hour of Proteinase K digestion and extensive washing, surface display of (PSMB7, Myl-6, DDX3X, EEF1E1, H2BC11) was restored in a time-dependent manner (n = 5). No recovery occurred when cells were incubated at 4 °C or treated with 350 nM brefeldin A (BFA). **(D)** Recovery of Inside-Out protein display after Heparinase II treatment. Similar to Proteinase K, surface expression of (PSMB7, Myl-6, DDX3X, EEF1E1, H2BC11) reappeared over time after Heparinase II removal, but not under conditions of 4 °C or 350 nM BFA (n = 5). **(E)** Stress-induced display of Inside-Out proteins in PBMCs. Untreated PBMCs exhibited no surface expression, but robust display was induced by 30 μM GW4869 for 1 hour (n = 40). Co-treatment with 350 nM BFA significantly inhibited this effect. (n = 40, *\*P ≤* 0.05, \**\*P ≤* 0.01, \*\**\*P ≤* 0.001, \*\*\**\*P ≤* 0.0001. Two-way ANOVA). Prolonged BFA incubation reduces Inside-Out protein display. A 24-hour treatment of HCT116 cells with 350 nM BFA markedly suppressed surface presentation, indicating a dynamic turnover of Inside-Out proteins at the membrane (n = 40, *\*P ≤* 0.05, \**\*P ≤* 0.01, \*\**\*P ≤* 0.001, \*\*\**\*P ≤* 0.0001. One-way ANOVA).

Subsequently, we examined the potential contribution of heparan sulfate to the cell surface attachment of I-O proteins. Heparan sulfate proteoglycans (HS) are well-established mediators of protein retention at the plasma membrane, facilitating protein clustering and cell signaling through interactions between their negatively charged sulfate and carboxyl groups and the positively charged regions on protein surfaces (40). These multivalent interactions generate robust, specific protein binding that surpasses the stability of interactions typically disrupted by high salt concentrations. Notably, treatment of HCT116 cells with Heparinase II, an enzyme that cleaves heparan sulfate chains, resulted in a significant decrease in I-O Fab binding, analogous to the effect observed with proteinase K (Fig. 4A). This finding underscores the pivotal role of heparan sulfate in anchoring I-O proteins to the surface of cancer cells, corroborating previous studies on HS-mediated protein association (41, 42). Furthermore, RNA-associated I-O proteins were also effectively removed by Heparinase II, indicating that heparan sulfate–mediated retention exceeds RNA-dependent mechanisms, thereby establishing a hierarchy of factors governing the surface presentation of I-O proteins.

To clarify the molecular basis of I-O protein interactions with heparan sulfate proteoglycans, we performed a heparin-agarose binding assay using nine I-O proteins with different surface charge profiles. All proteins showed strong, specific heparin binding, indicating high affinity for sulfated glycans. Supporting this, computational docking using ClusPro revealed distinct, positively charged surface pockets on each protein, characteristic of typical heparin-binding motifs (43) (Fig. S8). These findings suggest that the charge distribution of I-O proteins enables selective orientation and interaction with heparan sulfate, affecting which epitopes are exposed or occluded. This mechanism likely governs available protein-protein interactions at the cell surface, providing a rationale for why only a subset of Fabs that recognize the soluble protein also bind to the membrane-associated form, due to changes in epitope accessibility in the presence of HS.

Subsequently, we aimed to investigate the affinities of I-O proteins to the cell surface. This is presumably influenced by several factors related to the I-O protein’s HS association properties, such as the number, size, and distribution of their putative heparin-binding motifs. As will be elaborated in a subsequent section, the dynamics of recycling I-O proteins between the intracellular compartments and the cell surface further complicate this relationship. Nonetheless, we conducted this investigation using surfen (bis-2-methyl-4-amino-quinolyl-6-carbamide), a small molecule exhibiting an affinity of approximately 3 µM for HS at sites that compete with other potential binding entities, such as proteins (44). For this purpose, we selected eight I-O proteins with diverse sizes and surface characteristics. Notably, unlike Heparinase II, surfen is a non-destructive technique and can therefore be utilized in the determination of IC_50_ values within the context of the I-O) proteins. It is noteworthy that, although the shapes of the curves vary somewhat, all exhibited a similar affinity range between 70 and 250 nM (Fig. 4B).

After establishing that I-O proteins are strongly tethered to the plasma membrane through electrostatic interactions with HS, we examined whether the addition of high concentrations of soluble I-O antigens to surfen-treated cells would facilitate their adherence to the HS precleared surface. Using eight representative I-O antigens, both treated and untreated cells were incubated with 200 nM of the proteins for 30 minutes. The experiments were performed at 4°C to prevent the repopulation of antigens on the cell surface. Flow cytometry analysis revealed that there was essentially no difference between the untreated and precleared cells, both of which displayed no binding, indicating that simply adding high concentrations of soluble antigens does not facilitate their attachment to the plasma membrane, even when incubating them under nonphysiological conditions that should promote their binding (Fig. S9). This virtually eliminates the possibility that I-O protein display involves exogenous surface contamination.

### Trafficking of I-O Protein to the Cell Surface is a Dynamic Process

Despite the substantial removal of I-O proteins following surface digestion with proteinase K or Heparinase II, the treated HCT116 cells remained viable and exhibited rapid recovery. This observation suggests the reversible nature of I-O protein surface display. To examine whether the surface display of I-O proteins was replenished over time, as well as to elucidate the kinetics and specificity of this recovery process, we monitored the translocation of I-O proteins from intracellular pools to the cell surface over a 24-hour period employing the panel of 40 I-O-specific Fab antibodies. Both enzymatic treatments produced an immediate and significant loss of Fab binding across all targets (Fig. 4A). Follow-up time-course analyses were conducted at 37°C and 4°C.

At physiological temperature (37°C), substantial restoration of surface-displayed I-O proteins was evident as early as three hours post-digestion, with near-complete recovery by six hours, and display levels comparable to those of pre-digestion controls (Fig. 4C, D). Notably, recovery after Heparinase II treatment was particularly striking, given that reestablishment of the heparan sulfate layer is required to mediate I-O protein attachment. In contrast, following treatment with either enzyme, no recovery was observed at 4°C over the same interval, suggesting that I-O protein surface replenishment is an active, temperature-dependent process. To clarify the underlying trafficking mechanism, we inhibited ER-Golgi transport using Brefeldin A (BFA) (45) during surface digestion. In all conditions-regardless of the enzyme or incubation temperature-BFA completely prevented the restoration of surface I-O proteins (Fig. 4C, D). Consistently, BFA also blocked surface display of I-O proteins in GW4869-stressed PBMCs (Fig. 4E). Moreover, continuous treatment of HCT116 cells with BFA for 24 hours led to a nearly 60% reduction in I-O antigen surface levels (Fig. 4F), further supporting a critical requirement for ongoing ERGolgi trafficking in the maintenance and replenishment of I-O proteins at the plasma membrane.

### Dynamic Bidirectional Trafficking and Recycling of InsideOut

The restoration of inside-out proteins following their removal by Heparinase II and proteinase K exemplifies the dynamic nature of their surface presentation. To further investigate whether this trafficking is unidirectional or also involves internalization and recycling, we observed the internalization of surface-exposed I-O proteins over time. Utilizing the Incucyte S3 live-cell imaging system in conjunction with a pH-sensitive anti-IgG reporting system, where antibody internalization induces red fluorescence in acidic lysosomal compartments, we conducted experiments using PC3 and OVCAR3 cells, chosen for their size and suitability for quantitative imaging.

Our findings demonstrated a clear, time-dependent accumulation of red fluorescence within cells for numerous inside-out Fabs tested. Significantly, the internalization characteristics were enhanced when utilizing the IgG form of the antibody. The internalization of the IgG antibody targeting Myl6 is illustrated in Fig. 5A: there was no signal at baseline and substantial intracellular red fluorescence was observed after 18 hours (Fig. S10), confirming appreciable protein uptake. Quantitative analysis, conducted with imaging every 60 minutes, indicated a consistent increase in intracellular fluorescence, signifying dynamic internalization. Importantly, control cells incubated with an isotype IgG directed against showed no red fluorescence (Fig. 5A, S10), thereby verifying the specificity of the process.

**Figure 5.**
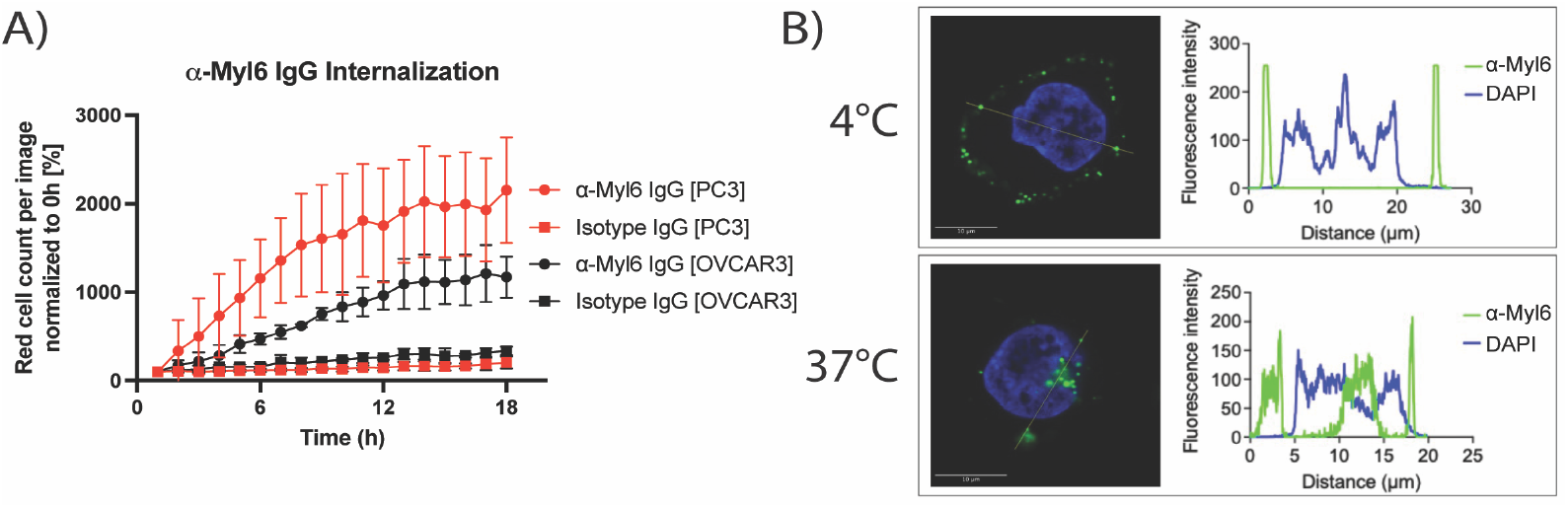
Internalization of Myl6 in cancer cells monitored by real-time live-cell imaging and confocal microscopy. **(A)** PC3 and OVCAR3 cells were subjected to real-time monitoring of anti-Myl6 IgG internalization using the Incucyte S3 live-cell analysis system. Quantitative analysis of red fluorescence over time reveals a continuous, time-dependent increase in anti-Myl6 IgG uptake, while the isotype control antibody exhibits no detectable internalization (n = 3, mean ± SD). **(B)** Confocal microscopy images of HCT116 live-cell immunostaining performed at 4°C or 37°C. An absence of anti-Myl6 IgG internalization at 4°C is demonstrated by fluorescence signal restricted to the cell surface, whereas prominent intracellular accumulation is observed at 37°C, confirming active, temperature-dependent uptake. Scale bar denotes 10 *μ*m.

To corroborate these observations in a different model and at a higher resolution, confocal microscopy was conducted on HCT116 cells. The cells were subjected to treatment with anti-Myl6 IgG at either 4°C or 37°C, followed by extensive washing, permeabilization, and staining with a secondary antibody. At 37°C, a marked internal signal was observed, indicative of active internalization, whereas no significant uptake was detected at 4°C (Fig. 5B), aligning with the temperature dependence of endocytosis and excluding permeabilization artifacts. Collectively, these findings provide strong evidence that the surface presentation of inside-out proteins in cancer cells is both dynamic and reversible. Rather than being irreversibly externalized, insideout proteins are capable of cycling back into cells, thereby demonstrating a continuous and time-dependent trafficking process.

## Discussion

This study establishes a new paradigm in protein trafficking by demonstrating that large cohorts of intracellular-origin (I-O) proteins can be dynamically displayed at the surface of cancer cells through a stress-regulated, heparan sulfate–dependent mechanism. Unlike canonical secretion pathways, which rely on signal peptides and vesicular transport, the display of I-O proteins represents a distinct mode of surface presentation tightly coupled to cellular stress and adaptive homeostasis. By leveraging APEX2-mediated proximity labeling and a comprehensive antibody-based screening strategy, we systematically mapped their surface occurrence, the kinetics of their surface residence and selectivity in tumor cells, revealing both the surprising diversity of the I-O surface proteome and its linkage to stress-responsive networks.

Our proteomic analysis identified approximately 140 high-confidence inside-out (I-O) proteins expressed on tumor surfaces. We noted that this I-O proteome is not comprehensive due to limitations in the biotinylation method used (46), which results in an underestimation of the total number of I-O proteins. Furthermore, the extent of surface expression of individual I-O proteins exhibits some variability among the cell lines examined; consequently, some proteins with lower expression levels may not attain the acceptance threshold in certain cell lines. However, while establishing a complete I-O proteome is a key future goal, this study was conducted to explore the characteristics and biological properties of the I-O protein phenomenon. Interestingly, the identified I-O proteins belong to families that include translation factors, ribosomal and proteasomal subunits, heat shock proteins, and other essential cell regulators. Gene ontology analysis revealed a strong tendency toward stress-related pathways, highlighting how I-O surface presentation aligns with broader adaptive response mechanisms.

To confirm and extend our proteomic findings, we generated a panel of nearly 500 antibodies targeting 40 selected I-O antigens, employing a high-throughput phage display platform and multiplexed assay system to enable comprehensive profiling of antibodies against all the selected targets. Surface abundance of these I-O proteins was validated across seven different cancer cell lines, with surface display levels varying somewhat by cell type (Fig. 2C). *In vivo* fluorescence imaging of a panel of six antibodies representing diverse families of I-O proteins showed that all the antibodies specifically accumulated in tumors within a breast cancer xenograft mouse model (Fig. 3A). The xenograft tissues exhibited profiles consistent with those observed in cultured cells, whereas normal kidney and liver tissues showed negligible binding. Importantly, I-O protein display was not detectable on peripheral blood mononuclear cells (PBMCs) under normal conditions (Fig. 2C) but became clearly visible after PBMCs were exposed to a stress-inducing inhibitor (Fig. S6). This stress-induced phenotype was reversible after inhibitor removal, highlighting the role of cellular stress in the surface display of I-O proteins.

As mentioned, I-O proteins are typically associated with families of proteins that play a role in cellular stress response activities. However, it’s still unclear whether they carry out any functions on the cell surface related to their natural intracellular activities. A substantial body of literature describes potential extracellular roles of members of the heat shock protein family, as well as other “moonlighting” proteins previously discussed. These extracellular functions are typically quite different from their roles inside the cell. The idea that each surface-displayed I-O protein or complex has a specific extracellular function seems unlikely, given the compartmentalized nature of intracellular environments, a feature that is missing on the cell surface. Therefore, we conclude that although some I-O proteins may have additional extracellular roles, overall, I-O proteins serve as a reservoir that can be efficiently shuttled between the inside and outside of the cell depending on stress conditions. This conclusion is supported by our observation that not all types of cellular stress affect the level of I-O proteins on the cell surface.

We further examined the molecular mechanisms behind I-O protein surface attachment. I-O proteins lack specific membrane localization signals or inherent modifications that directly anchor them to the plasma membrane. Proteinase K digestion removed most of the detectable surface I-O proteins, though the enzyme’s broad specificity limited insights into the exact mechanisms of attachment. In contrast, treatment of HCT116 cells with Heparinase II completely removed all 40 I-O proteins in the tested set (Fig. 4A). Additionally, incubating the cells with surfen, a small molecule inhibitor of heparan sulfate, non-destructively removed I-O proteins from the surface, and adding high concentrations of soluble I-O antigens afterward did not restore them (Fig. 4B, S10). This suggests that the surface-displayed I-O population on the cell surface is not attributable to capturing external cell debris. While other molecular components likely contribute to I-O protein attachment, the data indicate that heparan sulfate plays a key role in their membrane tethering.

This tethering of the I-O proteins via heparan sulfate mainly occurs through electrostatic interactions. The charge distributions and the various conformational states that heparan sulfate may assume in its cell-bound form imply that the presentation of the bound I-O proteins would be random. However, our data indicate a different picture: the orientations of the I-O proteins on the plasma membrane are not random. During the initial screening of Fab antibodies created through phage display selection targeting the soluble forms of I-O proteins, it was observed that only a subset of high-affinity Fabs for the soluble antigen also bound to cells. This indicates that certain epitopes are obscured by the membrane. Importantly, the same group of Fab molecules for nearly all I-O antigens consistently either binds or does not bind across various cell lines. We interpret this as the IO proteins being anchored in a way that ensures their antibody epitopes are consistently presented in similar orientations across all tested cell lines. This suggests that the heparan sulfate binding motifs on the surface of each I-O protein determine how they are oriented when attaching to the membrane surface.

A remarkable observation was the swift restoration of I-O protein display after enzymatic removal. When HCT116 cells were treated sequentially with proteinase K or Heparinase II at 37°C, surface levels of all 40 surveyed I-O proteins recovered to about 50% of their original levels within three hours and returned to pre-treatment levels by six hours. In contrast, recovery was nearly absent at 4°C or in the presence of the ER-Golgi trafficking inhibitor brefeldin A (Fig. 4C, D). This suggests that recycling of I-O proteins to the cell surface is dynamic, depends on temperature and ER function, and requires active intracellular trafficking.

Overall, these findings support a model where surface pools of I-O proteins are maintained in balance with their internal counterparts, regulated by cellular stress and facilitated through ER trafficking and heparan sulfate interactions. Furthermore, our antibody internalization experiments demonstrate that this process is bidirectional. It is observed that I-O proteins not only translocate from the interior of the cell to the surface but can also be recycled internally after reaching the exterior, likely through endocytosis, as this process does not occur at 4°C. The ultimate fate of these recycled proteins inside the cell remains unknown. An example of this process is how I-O proteins appear on stress-induced PBMCs and then disappear, probably through endocytosis when the stress subsides.

Quantifying the relative abundance of individual I-O proteins using flow cytometry reveals variation among different cell lines. However, within a specific cell line, the levels remain consistently similar across duplicated experiments. This suggests that the export and import mechanisms are nearly in balance. Notably, after treatment with either proteinase K or Heparinase II, the recovery levels of surface display across the tested targets generally maintain a consistent ratio at each time point. This implies that the mechanism controlling internal factors responsible for the abundance of surface display I-O proteins under stress conditions is activated early in the process. An important future objective is to ascertain the population of the I-O proteins within and outside the cell, as well as to analyze how this varies among different proteins and cell lines. This represents a substantial undertaking and is therefore beyond the scope of this study.

A particularly striking finding was the strong compositional overlap between surface I-O proteins and stress granule (SG) components, with nearly half of detected I-O proteins also annotated in SG databases (47) (Fig.1C). This convergence reinforces the close connection between stress adaptation pathways operating in distinct compartments of the cell. Yet, important differences emerged: whereas SGs are highly ordered, cytoplasmic condensates that require external stress triggers for assembly (48–50), I-O surface display occurs broadly in unstimulated cancer cells and involves folded proteins bound through extracellular heparan sulfate, rather than structured, condensate-like assemblies. More recently, growing evidence suggests that SGs are not just storage units but are organized to perform advanced intragranular functions (51). When cell stress subsides, SGs begin to dissolve and release their components for cellular use (52). The stress reduction also affects the I-O surface display phenomenon by lowering display levels; however, it remains unclear whether any of the affected I-O proteins are ultimately integrated into intracellular networks. These differences suggest that while SGs and I-O proteins draw from overlapping functional pools, they represent distinct arms of the stress response-SGs providing an adaptive, compartmentalized strategy and I-O surface proteins offering a rapid, membrane-associated mechanism for maintaining homeostasis. Furthermore, it is probable that there is turnover within the I-O protein pool, characterized by a dynamic equilibrium between expression and degradation, depending on the stress level.

The discovery that non-secretory proteins can be stably displayed on the tumor cell surface through stress-induced, heparan sulfate–mediated mechanisms has broad biological and translational implications. Functionally, surface I-O proteins may enable cancer cells to rapidly adjust their interactions with the microenvironment, potentially influencing nutrient uptake, signal communication, or immune detection during stress fluctuations. The observed process of recovery and recycling of I-O display after removal suggests that this pathway is actively maintained rather than incidental, indicating it may be a stress adaptation mechanism of previously unrecognized significance. Clinically, the tumor-specific surface localization of I-O antigens verified in xenograft models underscores their potential as accessible biomarkers and therapeutic targets. Antibody-based strategies targeting I-O proteins could exploit their selective expression in tumors and their inducibility under stress, offering a new opportunity for cancer treatment.

Overall, these findings turn intracellular protein surface display from a rare anomaly into a specific biological pathway, regulated at the intersection of stress response, ER trafficking, and heparan sulfate-mediated tethering. The similarities with stress granule biology suggest a coordinated system for quick, adaptive stress responses, occurring both inside the cell and on its surface. Besides answering key questions about protein localization, this work uncovers a new layer of proteome dynamics with important implications for cancer biology, immune surveillance, and therapeutic strategies. Future studies focusing on the molecular signals that control trafficking and the functional consequences of surface I-O localization will be essential for understanding how stress responses are integrated into tumor biology and how they can be targeted for therapy.

## Supporting information

I-O_Slezak_Supplement

## Acknowledgements

We thank the members of the Kossiakoff Laboratory for their support, with special appreciation to Carla Meints and Tomasz Zamyslowski. We are grateful to Theodore Steck and David Pincus for their valuable suggestions. We also acknowledge the University of Chicago Sanger DNA Sequencing Core for sequencing services and the University of Chicago Integrated Light Microscopy Core (RRID: SCR_019197) for confocal imaging. We extend our appreciation to the staff at the University of Chicago Proteomics Platform. Finally, we gratefully acknowledge the support provided by the Chicago Biomedical Consortium and the University of Chicago Comprehensive Cancer Center.

## Author contributions

T.S, K.M.O, and A.A.K designed research; T.S generated new reagents; T.S, K.M.O, T.G.A, J.L, N.M, A.A, E.F.S, DA.L. performed research; T.S analyzed data; T.S, K.M.O and A.A.K wrote the paper.

## Materials and Methods

### APEX2 Proximity Labeling

HCT116 cells were cultured to ∼90% confluency. Following trypsinization and two phosphate buffered saline (PBS) washes, cells (6 × 10^6^ per replicate) were resuspended in 1 mL 200 nM APEX2 solution. After the addition of 1 mL substrate solution (50 µg/mL biotin-PEG4-phenol, 0.1% H_2_O_2_ in PBS) and incubation for 60 seconds, 1 mL quencher solution (50 mM NaN_3_, 100 mM sodium ascorbate, 50 mM Trolox in PBS) was added. Cells were washed twice in PBS, then incubated with 50 mM Proteinase K (Invitrogen) in reaction buffer (50 mM Tris-HCl, 7.5 mM CaCl_2_, pH 7.5; 1 mL per 4 × 10^6^ cells) or reaction buffer alone (control) for 1 hour on ice with occasional mixing. Subsequent washes in reaction buffer removed the enzyme.

### Mass Spectrometry

Cells were lysed in 400 µL RIPA buffer (Millipore Sigma) on ice for 30 minutes with occasional mixing. Lysates were clarified by centrifugation, and protein concentrations were determined using the Bradford assay (Thermo Scientific). For each sample, 250 µg lysate was applied to columns containing 400 µL NeutrAvidin Agarose Resin (Thermo Scientific), equilibrated in RIPA. Following a 3-hour incubation at 4°C with rotation, columns were washed with 10 mL each of RIPA, PBS/1M NaCl, and 50 mM NH_4_HCO_3_/2 M urea. Columns were subjected to on-resin reduction and alkylation with 100 µL LYSE solution (iST Sample Preparation Kit, PreOmics) at 55°C, 1000 rpm for 10 minutes, followed by digestion with 50 µL DIGESTION solution at 37°C, 500 rpm for 90 minutes. Supernatants were collected, 100 µL STOP solution added, and samples desalted and eluted per PreOmics protocol. Peptides were vacuum-dried and resuspended in LCLOAD solution for LC-MS/MS. Mass spectrometry was performed on an Exploris 480 (Thermo Scientific) coupled to an UltiMate 3000 nanoLC system. Peptides were separated on a 50 cm MonoCap column (0.75 mm ID, GL Sciences) at 500 nL/min, 25°C, with the following solvent B (0.1% formic acid in acetonitrile) gradient: 0–5 min: 5%; 5–105 min: 22%; 105–124 min: 34%; 124–125 min: 95%; 125–130 min: 95%; 130–131 min: 4%; 131–145 min: 4%. Full scan MS spectra were acquired from m/z 350–1200 at 120,000 resolution, 50 ms maximum injection time, and AGC target 3e6. The cycle time was 3 s; the intensity threshold was 5e4. MS/MS scans used 15,000 resolution, 40 ms maximum injection time, AGC target 4e4, isolation window 1.6 m/z, first mass 110 m/z, and NCE 30. Dynamic exclusion was set to 30 seconds and charge states 2–7 were analyzed. Raw data were analyzed using Proteome Discoverer v3.0.0.757 (ThermoFisher), searching against the human protein database with the Sequest HT engine. Mass tolerances were set to 10 ppm (precursor) and 0.02 Da (fragment). Peptides were filtered at a 1% FDR using the Target Decoy PSM Validator. Trypsin was specified as the protease (full specificity; up to three missed cleavages; minimum peptide length 7 aa, maximum 35 aa). Fixed modification: carbamidomethylation (+57.021 Da, cysteine); dynamic modifications: oxidation (+15.995 Da, methionine), protein N-terminal methionine loss (−131.040 Da), N-terminal acetylation (+42.011 Da), N-terminal methionine loss with acetylation (−89.030 Da). Protein abundance tables were exported to Perseus (v2.1.1.0, MaxQuant) (53). Statistical significance was determined using two-sample t-tests. Protein abundance fold changes were calculated as the ratio of the median abundance in non-proteinase K-treated cells to that in treated cells. Enrichment and network analyses were performed with STRING v12.0 and Cytoscape v3.10.2 (54, 55). Visualizations were generated in GraphPad Prism and Origin 2025 v10.2.0.196. The mass spectrometry proteomics data have been deposited to the ProteomeXchange Consortium via the PRIDE (56) partner repository with the dataset identifier PXD069264.

### Protein Cloning, Expression and Purification

Target proteins were cloned into the pHFT2 vector. An N-terminal Avitag and His-tag were incorporated to facilitate efficient purification and in vitro biotinylation. Proteins were expressed in *E. coli* BL21 (DE3) cells. Expression was induced at an OD600 of 0.6 with 1 mM IPTG, followed by overnight incubation at 20°C. Cells were harvested and lysed by sonication in buffer A (50 mM Tris-HCl, pH 8.0; 150 mM NaCl; 10% glycerol). Lysates were clarified by centrifugation and applied to TALON resin (Takara Bio) for immobilized metal affinity chromatography (IMAC). Bound proteins were eluted with buffer A supplemented with 150 mM imidazole. For several targets, purification was performed from inclusion bodies. Inclusion bodies were solubilized in 6 M guanidine hydrochloride (Gua-HCl) in buffer A containing 0.3 mM TCEP. Denatured proteins were purified on TALON resin and subjected to on-column refolding by gradual buffer exchange consisting of six sequential two-fold Gua-HCl dilutions, followed by a final wash in buffer A. Proteins were eluted as above. Protein biotinylation was performed enzymatically using Avi-tagged proteins and the BirA ligase, following the manufacturer’s protocol (Avidity).

Selected Fabs were subcloned from phage display outputs into the RH2.2 expression vector. Fabs were expressed in the periplasm of *E. coli* BL21 cells at 37°C for 4 hours post-induction (1 mM IPTG, OD600 = 0.8–1.0). Cells were harvested and lysed by sonication in Protein G wash buffer (50 mM sodium phosphate, 500 mM NaCl, pH 7.4), and the lysate was clarified by centrifugation. The Fab-containing supernatant was loaded onto a Protein G-F affinity column (57). Elution was performed with 0.1 M glycine, pH 2.6, and immediately neutralized with 1 M Tris-HCl, pH 8.5. Fabs were dialyzed overnight into HBS buffer. For conversion into full-length IgG, Fab sequences were reformatted with the human IgG1 Fc domain in the pSCSTa vector. An N31D mutation was introduced to abrogate an N-linked glycosylation site adjacent to CDR-H2. Plasmids encoding heavy and kappa light chains (7.5 µg of each, pSCSTa backbone) were transiently transfected into Expi293 cells using Opti-MEM medium (Gibco) and the ExpiFectamine 293 kit (Gibco) per manufacturer’s instructions. After five days, supernatants were collected, and IgG was purified as above using Protein GF affinity chromatography (57), eluted with 0.1 M glycine (pH 2.6), neutralized, and dialyzed into HBS.

### Phage Display Selection Protocol

High-affinity binders were obtained through five rounds of phage display selection, using protocols adapted from previously published methods (32, 33). Briefly, biotinylated target proteins were immobilized on streptavidin-coated paramagnetic beads (Promega), followed by incubation with the phage display library (1 × 10^10^ cfu) for 1 hour at room temperature with gentle agitation. In the first round, 1 µM target protein was added to 200 µL beads. Unbound phages were removed by three washes, and beads were added to log-phase *E. coli* XL-1 Blue cells (Stratagene) and incubated for 20 minutes at room temperature. Phage were amplified overnight at 37°C in medium containing 100 µg/mL ampicillin and 1 × 10^9^ pfu/mL M13K07 helper phage (NEB). Amplified phages were precipitated using 20% PEG/2.5 M NaCl on ice for 20 minutes. Before each selection round, phage pools underwent negative selection with empty beads for 30 minutes to deplete non-specific binders. In subsequent rounds, target antigen concentration was sequentially decreased (1 µM, 200 nM, 50 nM, 10 nM, 1 nM; rounds 1–5, respectively). After selection, beads were washed five times with PBST containing 0.5% bovine serum albumin (BSA). Bound phages were eluted with 0.1 M glycine (pH 2.6), neutralized with Tris-HCl (pH 8.0), and used for infection and amplification as above. In rounds 4 and 5, *E. coli* cultures were plated on ampicillin agar, 192 colonies were screened by single-point phage ELISA to identify high specificity clones, which were sequenced and subcloned into RH2.2 for further characterization.

### Phage Enzyme-Linked Immunosorbent Assay (ELISA)

For phage ELISA, target proteins (50 nM) were immobilized in high-binding microplate wells (Greiner) for 30 minutes. Plates were blocked with 2% BSA for 1 hour at room temperature. After 15 minutes of phage incubation, wells were washed three times with 0.5% BSA/PBST and then incubated with HRP-conjugated Protein L (Thermo Scientific; 1:5000 in HBST) for 20 minutes. Plates were developed with TMB (Thermo Scientific), the reaction was quenched with 10% H_3_PO_4_, and absorbance was measured at 450 nm.

### Multipoint ELISA

Target proteins (50 nM) were immobilized in high-binding wells, blocked as above, and subjected to eight serial three-fold dilutions for each construct. After 20 minutes’ incubation, wells were washed, then incubated with HRP-conjugated anti-human (Fab)_2_ antibody (Jackson ImmunoResearch, 1:5000 in PBST) for 20 minutes. Plates were developed with TMB, quenched with H_3_PO_4_, and absorbance was read at 450 nm. The EC50 values were calculated using GraphPad Prism.

### Flow cytometry

Cancer cell lines (ATCC) were cultured in appropriate media supplemented with 10% FBS and 1% penicillin/streptomycin. PBMCs were isolated from healthy individuals and stored frozen until use. Cells were stained with 200 nM Fab for 30 minutes on ice, washed three times with PBS containing 1% BSA, and detected using AffiniPure Goat Anti-Human IgG F(ab’)_2_ (Jackson ImmunoResearch; 1:400 in PBS/1% BSA) for 20 minutes on ice. Following three further PBS/1% BSA washes, data were acquired on a CytoFLEX flow cytometer (Beckman Coulter) and analyzed using FlowJo 10.10.0 (BD Biosciences).

### Confocal Microscopy

HCT116 cells were seeded at ∼50,000 cells/mL (300 µL/well) on 8-well chamber slides (ibidi) and cultured for 2 days in McCoy’s 5A medium (ATCC) supplemented with 10% FBS (Gibco) at 37°C and 5% CO_2_. Cells were incubated with 200 nM Fab in complete medium at 37°C for 15 minutes, followed by one PBS wash. Secondary detection was performed with Alexa Fluor 647-conjugated AffiniPure™ Goat Anti-Human IgG, F(ab’)_2_ (Jackson ImmunoResearch, 1:200 in PBS/1% BSA) at 4°C for 20 minutes and subsequently washed twice. Cells were fixed with 4% paraformaldehyde for 10 minutes, stained with 1:1000 DAPI (ThermoFisher) in PBS/1% BSA for 5 minutes, and washed three times. The cells were imaged using a Leica SP8 confocal microscope and the image analysis was conducted in Fiji/ImageJ.

### Internalization Validation by Confocal Microscopy

Cells were incubated with 200 nM I-O IgG in complete media at 37°C for 15 minutes, washed once with PBS, then incubated with AffiniPure™ Goat Anti-Human IgG, F(ab’)_2_ (1:200, Jackson ImmunoResearch) in PBS/1% BSA at 37°C for 20 minutes. After switching to complete media, cells were incubated for 3 additional hours at 37°C. Cells were then washed, fixed, and stained with DAPI as described above. Imaging and analysis were performed as above.

### Live cell internalization analysis

PC3 cells were seeded at a density of 10,000 cells per well in black-walled 96-well plates (Thermo Scientific) and incubated overnight to allow for adherence. α-Myl6 IgG and isotype control IgG were employed at a final concentration of 4 µg/mL and conjugated with the Incucyte Human Fab-fluor-pH Antibody Labeling Dye for Antibody Internalization (Red) (Sartorius) according to the manufacturer’s protocol. Antibody internalization and cellular dynamics were monitored by acquiring images every hour using the Incucyte S3 Live-Cell Analysis System (Sartorius).

### Flow Cytometry After Treatments with Surface Modulators

HCT116 cells were cultured under standard conditions until approximately 90% confluency was achieved. For surface modulation studies, treatments were performed either on adherent cells or following cell suspension, depending on the specific reagent. NaCl and PNGase F treatments were conducted while cells remained adherent on the culture plate. For NaCl treatment, cells were washed with phosphate-buffered saline (PBS), incubated with 500 mM NaCl for 1 minute, washed again with PBS, and subsequently detached by trypsinization. For PNGase F treatment, cells were incubated directly on the plate with 50 U/mL PNGase F (New England Biolabs) in reaction buffer (50 mM Tris-HCl, 7.5 mM CaCl_2_, pH 7.5) at 37°C for 1 hour, followed by trypsinization to detach the cells. In contrast, treatments with RNase A, heparinase II, and proteinase K were conducted after cells were lifted with trypsin and resuspended. Detached cells were incubated in reaction buffer under the following conditions: RNase A at 20 µg/mL for 1 hour at 37°C; heparinase II (*Bacteroides eggerthii*, New England Biolabs) at 2 U/mL for 15 minutes at 37°C; and proteinase K (Invitrogen) at 500 µg/mL for 1 hour at 4°C. After the respective surface modifications, all cells were thoroughly washed and resuspended in PBS containing 1% bovine serum albumin (BSA). Cells were then seeded in 96-well plates at a density of 1 × 10^5^ cells per well. For immunostaining, cells were incubated with 200 nM antigen-specific Fab fragments for 20 minutes on ice, washed, and subsequently stained with Alexa Fluor 647–conjugated antihuman IgG F(ab’)_2_ secondary antibody (Jackson ImmunoResearch). After a final wash, samples were analyzed using a CytoFLEX flow cytometer (Beckman Coulter).

### PBMC Stimulation with GW4869, Sodium Arsenite and Torin-1

PBMCs from healthy donors were thawed, washed twice in RPMI 1640 (Corning) to remove DMSO, and incubated in complete medium with 30 µM GW4869 (MedChemExpress), 0.5 mM sodium arsenite (MilliporeSigma), 250 nM Torin 1 (MedChemExpress), or DMSO control in T-75 flasks at 37°C/5% CO_2_ for 1 hour. Half of the cells treated with 30 µM GW4869 were cultured for an additional 24 hours before analysis. Following incubation and two PBS washes, cells were analyzed immediately by flow cytometry as described above.

### Heparin-Agarose Affinity Chromatography

Heparin-Agarose (Sigma-Aldrich) resin was equilibrated with a solution of 10 mM Tris pH 8.0 containing 100 mM NaCl. Recombinant I-O proteins (100 μg) were incubated with the equilibrated resin for two hours at 4°C. Resin was washed extensively with a solution of 10 mM Tris pH 8.0 before elution using a solution of 10 mM Tris pH 8.0 containing 1.5 M NaCl. Affinity chromatography was assessed using sodium dodecyl sulfate polyacrylamide gel electrophoresis (SDS-PAGE).

### Flow cytometry after treatment with Surfen

HCT116 cells were cultured to 90% confluency, lifted with trypsin, and washed once in ice-cold PBS containing 1% bovine serum albumin (BSA). Cells were incubated with twofold serial dilutions of Surfen (Tocris) in PBS/1% BSA for 10 minutes on ice. Antibodies were added directly to a final concentration of 100 nM to cells treated with Surfen for an additional 20 minutes on ice. Cells were washed twice in cold PBS containing 1% BSA. Detection was performed using a secondary anti-human Fab’_2_ Alexa 647 antibody (Jackson ImmunoResearch; 1:400 in PBS containing 1% BSA) for 20 minutes on ice. Cells were washed three times in cold PBS containing 1% BSA and analyzed by flow cytometry. For flow cytometry analysis of biotinylated I-O proteins on Surfen-treated cells, HCT116 cells were cultured and harvested as described above. Cells were treated with 2 μM Surfen diluted in PBS/1% BSA or with PBS/1% BSA alone for 10 minutes on ice and then washed two times in PBS/1% BSA. Biotinylated I-O proteins were diluted to 200 nM in PBS/1% BSA and incubated with cells for 30 minutes on ice and then washed two times. Alexa Fluor 488 conjugated streptavidin (Invitrogen) was diluted to 2.5 μg/mL in PBS/1% BSA and incubated with cells for 20 minutes on ice. Cells were washed three times and then analyzed by flow cytometry.

### Recovery after Proteinase K treatment

HCT116 cells were treated with Proteinase K or control (buffer alone) for 1 hour on ice. Cells were washed twice with complete media to inactivate the enzyme. A portion was analyzed by flow cytometry immediately; the remaining cells were recovered in complete medium for 24 hours at either 37°C/5% CO_2_ (with or without 350 nM Brefeldin A in DMSO) or at 4°C. Flow cytometry was performed at 0.5, 1, 3, 6, 12, and 24 hours, with trypsinization of adherent cells required from 6 hours onward. Staining and analysis were performed as previously described.

### Recovery after Heparinase II treatment

HCT116 cells were treated with Heparinase II as described above. After enzyme inactivation, cells were either analyzed immediately or recovered for 24 hours in complete medium (with or without 350 nM Brefeldin A in DMSO or at 4°C). Time courses and flow cytometry analysis were identical to Proteinase K recovery experiments.

### In vivo I-O Fab accumulation in tumor xenografts

NOD.Cg-*Prkdc*^*scid*^ *Il2rg*^*tm1Wjl*^/SzJ (NSG) mice, sourced from The Jackson Laboratory, were maintained under specific pathogen-free (SPF) conditions with ad libitum access to water and food. All animal experiments were performed in strict accordance with Animal Care and Use Protocol (ACUP) 72785, as approved by the Institutional Animal Care and Use Committee of the University of Chicago. To generate the in vivo xenograft model, 6to 7week-old male NSG mice were subcutaneously inoculated with 5 × 10^6^ PC3 cells per mouse. After three weeks, once tumor xenografts reached a mean volume of approximately 500 mm^3^, mice were randomized into experimental groups (n = 2 per group). For the treatment arm, Fabs^LRT^ scaffold targeting I-O antigens (16 µM) were incubated with engineered protein GA1-Cy7 (38) (16 µM) at a 1:1 volume ratio for 10 minutes at room temperature. Subsequently, PC-3 xenograft-bearing mice received a single intravenous injection of 200 µL of the Fab^LRT^/GA1-Cy7 complex. Control mice were administered 200 µL of 8 µM GA1-Cy7 alone. At specified time points post-injection, animals were anesthetized with isoflurane (Dechra) and imaged using the Lago X in vivo imaging system (Spectral Instruments Imaging) to assess fluorescence accumulation within xenografts. Fluorescence intensity in tumor tissues was quantified using Aura software (Spectral Instruments Imaging).

### Flow cytometry staining using healthy murine kidney cells

Eight-week-old male NSG mice were euthanized via CO_2_ inhalation and subsequently dissected on ice. Kidneys were harvested and subjected to mechanical dissociation in Hanks’ Balanced Salt Solution (HBSS) using a scalpel. The resulting cell suspension was passed through a 70 µm cell strainer (Corning) and incubated in ACK (Ammonium-Chloride-Potassium) Lysing Buffer (Thermo Fisher) for 2 minutes. Cells were then washed with 2% FBS/PBS (Thermo Fisher), stained with 500 nM Fabs targeting I-O antigens at room temperature for 30 minutes, and subsequently incubated on ice for 30 minutes with DyLight™ 405 AffiniPure® Goat Anti-Human IgG, F(ab’)_2_ fragment specific (1:200 dilution; Jackson ImmunoResearch Laboratories). After rinsing with PBS, cells were stained on ice for 30 minutes with eBioscience™ Fixable Viability Dye eFluor™ 520 (1:1000 dilution; Thermo Fisher). The final cell preparations were rinsed with PBS and analyzed using an Aurora Flow Cytometer (Cytek Biosciences).

