## Supplementary material for "Dynamic translocation of Inside-Out proteins to the cell surface underlies cellular adaptation to cancer-induced stress": I-O_Slezak_Supplement

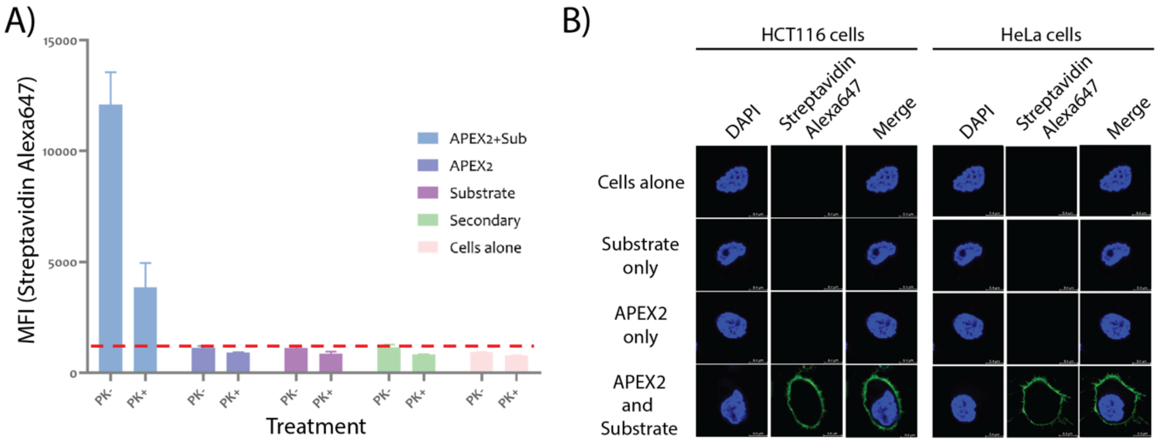


**Fig. S1. Validation of APEX2-mediated surface biotinylation in HCT116 and HeLa cells.** (A) Quantification of surface biotinylation efficiency. Cells were subjected to APEX2-mediated biotinylation followed by Proteinase K treatment, which selectively digests surface-exposed proteins. Flow cytometry analysis showed reduced Alexa Fluor 647-conjugated streptavidin signal post-treatment, indicating successful removal of biotinylated "Inside-Out" proteins, while residual signal confirms biotinylation of native membrane proteins. (B) Confocal microscopy images demonstrating robust surface-localized biotinylation in untreated HCT116 and HeLa cells, visualized via streptavidin-Alexa647 staining. Scale bar denotes 10 $\mu$m.

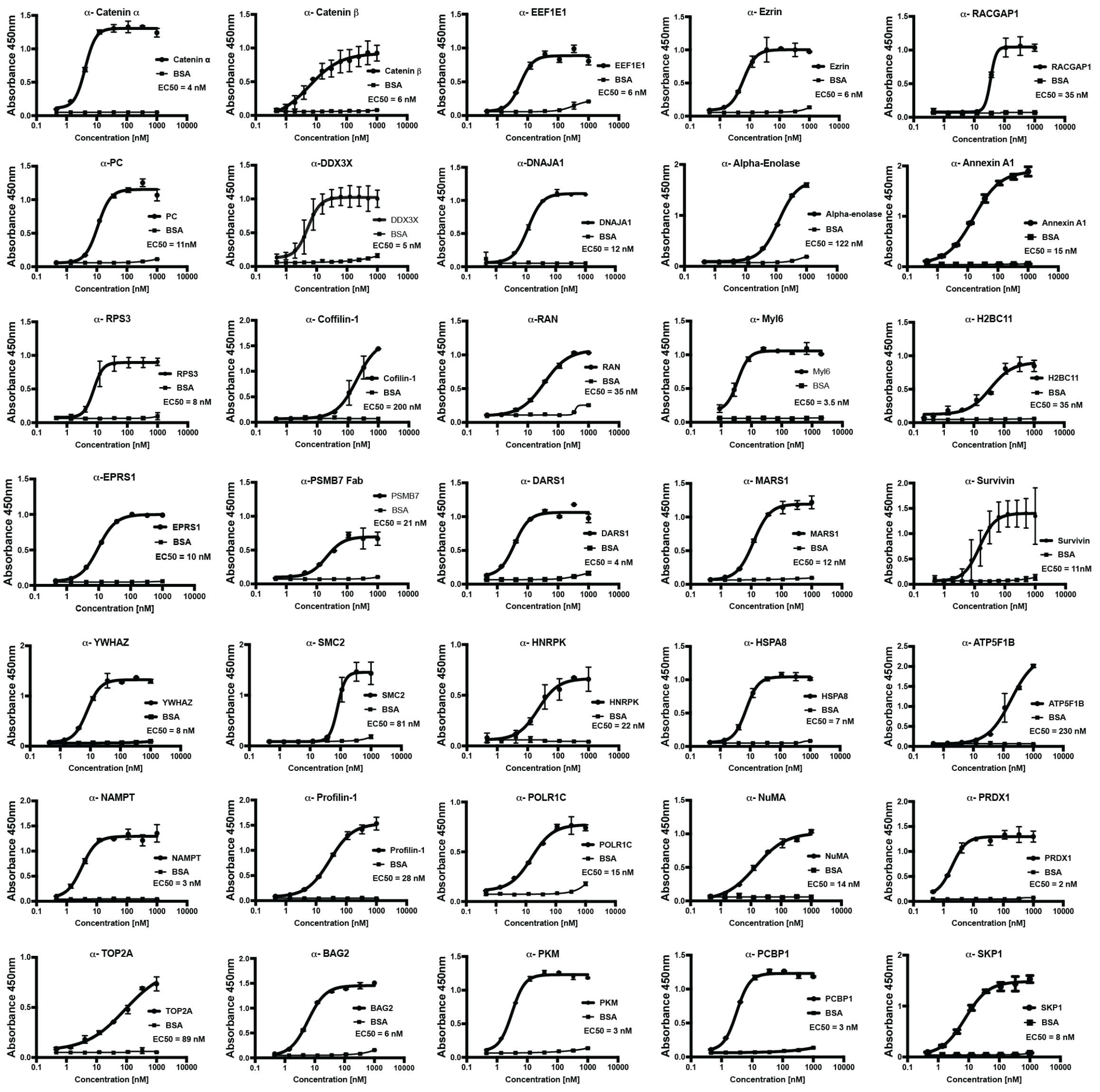


**Fig. S2.** Multi-point ELISA results are shown for representative antibodies generated via phage display against inside-out proteins. In total, 496 antibodies were engineered to target 40 distinct inside-out proteins.

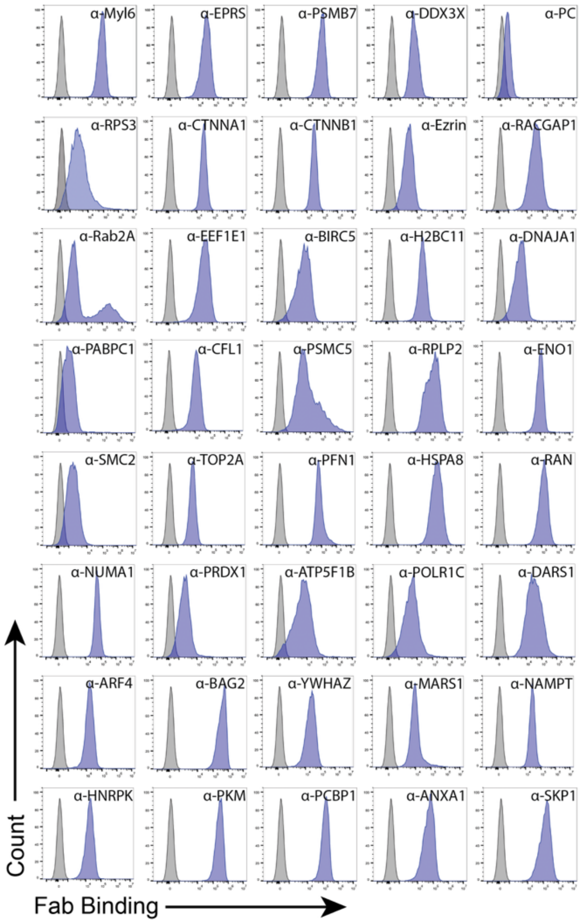


**Fig. S3.** Representative flow cytometry profiles demonstrating surface staining of the HCT116 cancer cell line with antibodies against Inside-Out proteins.

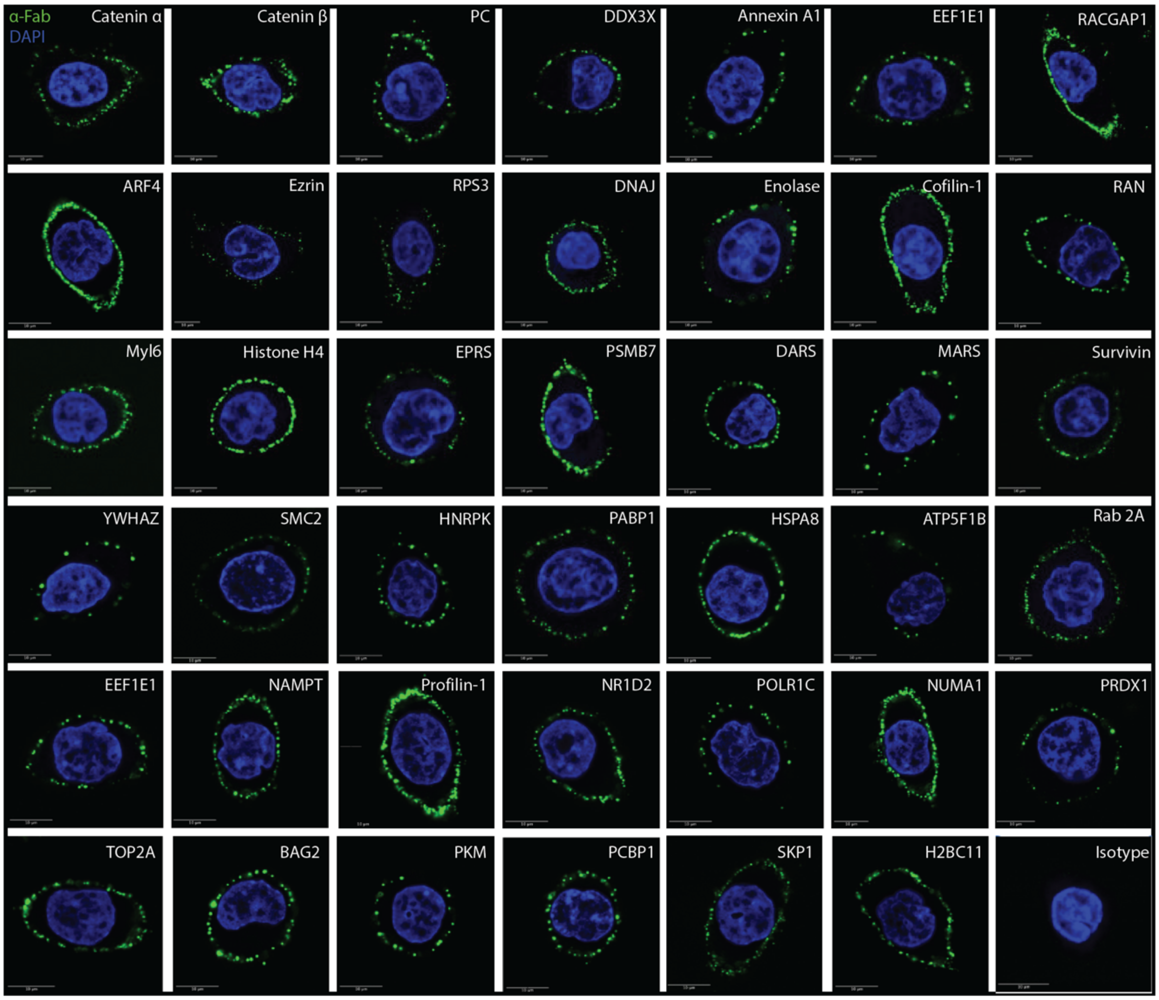


**Fig. S4. Cell surface immunofluorescence staining using antibodies against inside-out proteins.** Representative confocal microscopy results illustrate live-cell HCT116 surface staining with antibodies targeting inside-out proteins (green) and DAPI (blue). Scale bar denotes 10 $\mu$m.

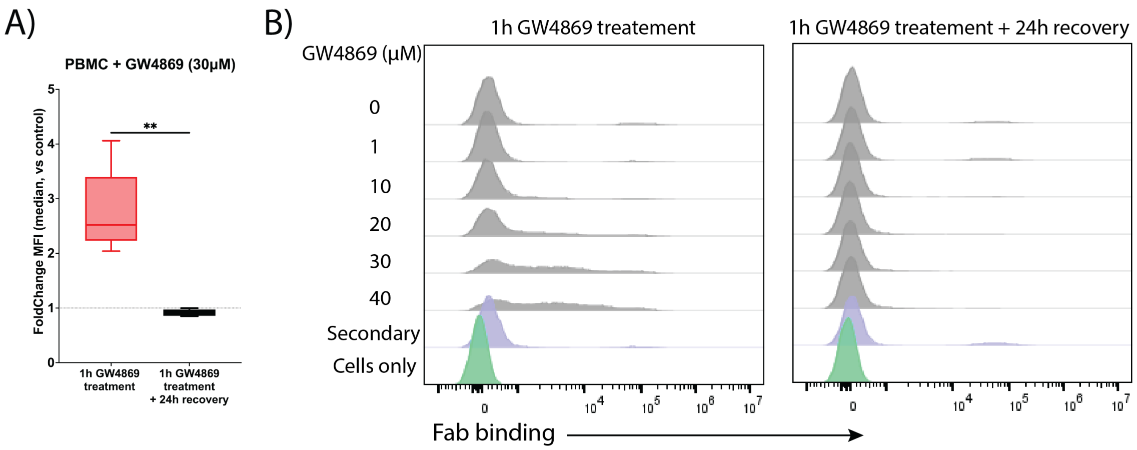


**Fig. S5. Stress-induced surface display of I-O proteins in PBMCs following GW4869 treatment.** (**A**) Box plot illustrating the fold change of the anti-I-O Fabs binding to PBMCs after a 1-hour GW4869 treatment, demonstrating the robust cell surface expression of I-O proteins (n = 6, mean ± SD. **P* ≤ 0.05, ***P* ≤ 0.01, ****P* ≤ 0.001, *****P* ≤ 0.0001. t-test). No binding is observed after a 24-hour recovery period post GW4869 removal. (**B**) Representative flow cytometry histograms showing concentration-dependent effect of 1-hour GW4869 treatment on the surface display of RPS3 in PBMCs and the absence of anti-RPS3 Fab binding to the cell surface following 24-hour recovery.

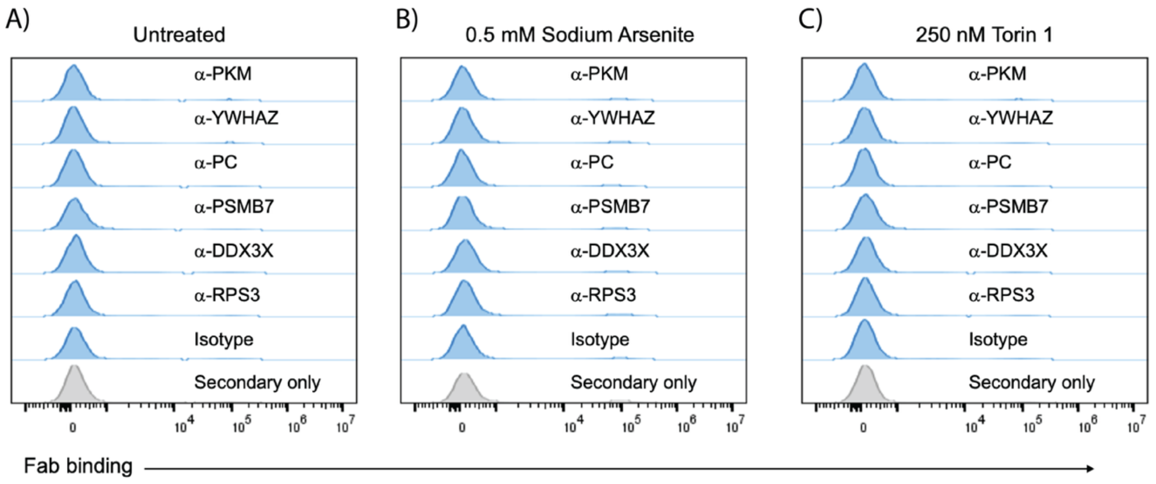


**Fig. S6. Flow cytometry analysis of I-O display in PBMCs after stress induction with Sodium Arsenite and Torin 1.** (**A**) Histograms for untreated PBMCs stained with the indicated antibodies. (**B**) Histograms for PBMCs treated with 0.5 mM sodium arsenite for 1 hour and then stained with the indicated antibodies. (**C**) Histograms for PBMCs treated with 250 nM Torin 1 for 1 hour and then stained with the indicated antibodies.

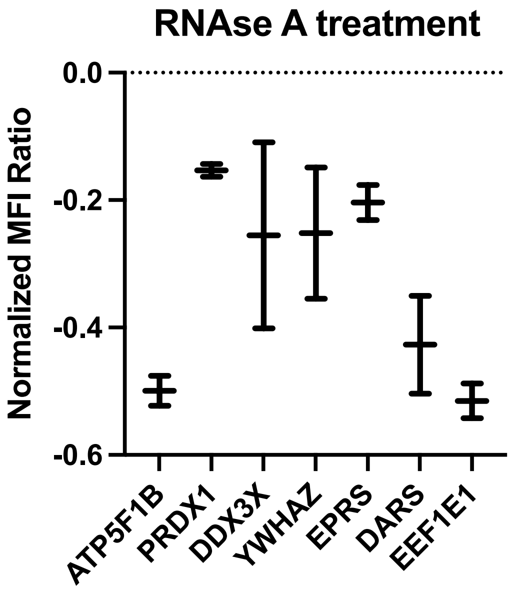


**Fig. S7. RNase A treatment selectively reduces the cell surface presentation of RNA-binding proteins on HCT116 cells.** HCT116 cells were treated with RNase A for 1 hour, resulting in reduced surface expression of various RNA-binding proteins. A moderate decrease in surface levels was observed for PRDX-1, DDX3X, YWHAZ, and EPRS, while ATP5F1B, DARS, and EEF1E1 exhibited a more pronounced reduction upon RNase A treatment. Quantification of binding was normalized to Fab binding specific to each target protein prior to RNase A incubation.

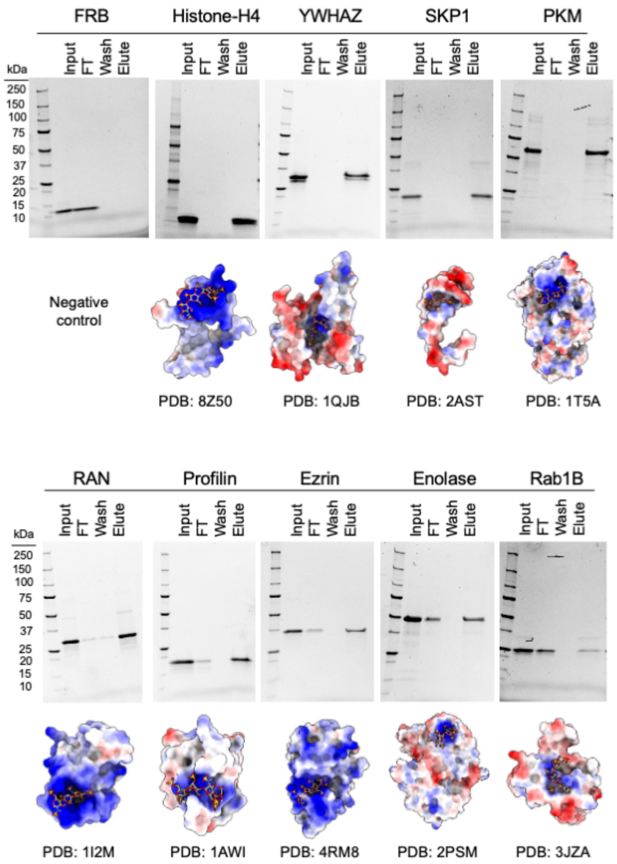


**Fig. S8. Heparin-agarose binding assay reveals binding of selected Inside-Out proteins to heparin.** Purified proteins were incubated with heparin-agarose resin and the samples from flow-through, wash, and elution fractions were analyzed by SDS-PAGE. The results confirm the interaction of Inside-Out proteins with heparin, but not the negative control (FRB). Below each gel, schematic models illustrate the predicted heparin-binding site for each protein using the ClusPro server.

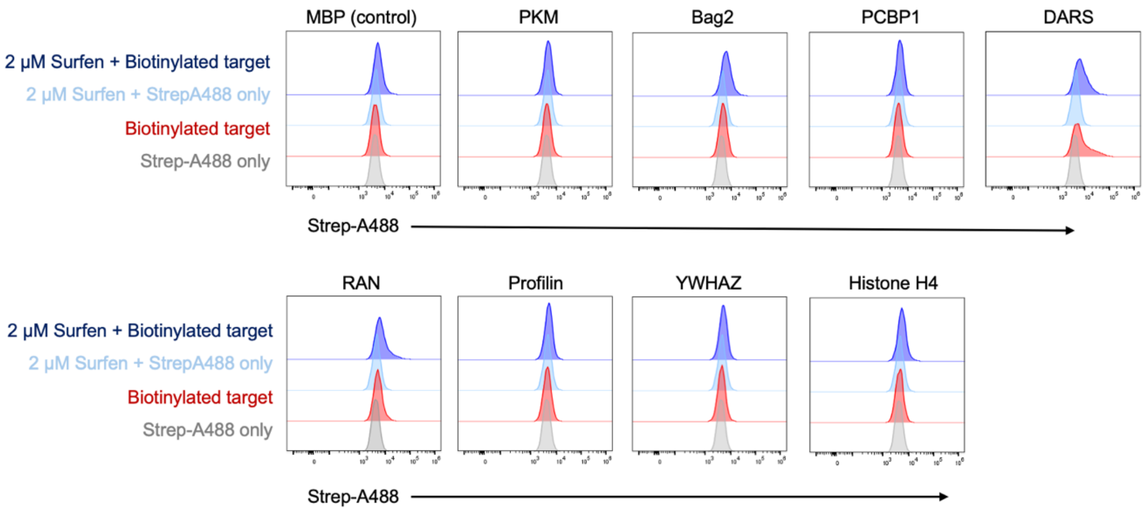


**Fig. S9. Flow cytometry analysis of biotinylated I-O antigens incubated with Surfen-treated HCT116 cells.** HCT116 cells were treated with or without 2 µM Surfen before incubation with the indicated soluble biotinylated I-O antigens for 30 minutes on ice. Secondary detection was performed using Alexa Fluor 488-conjugated streptavidin.

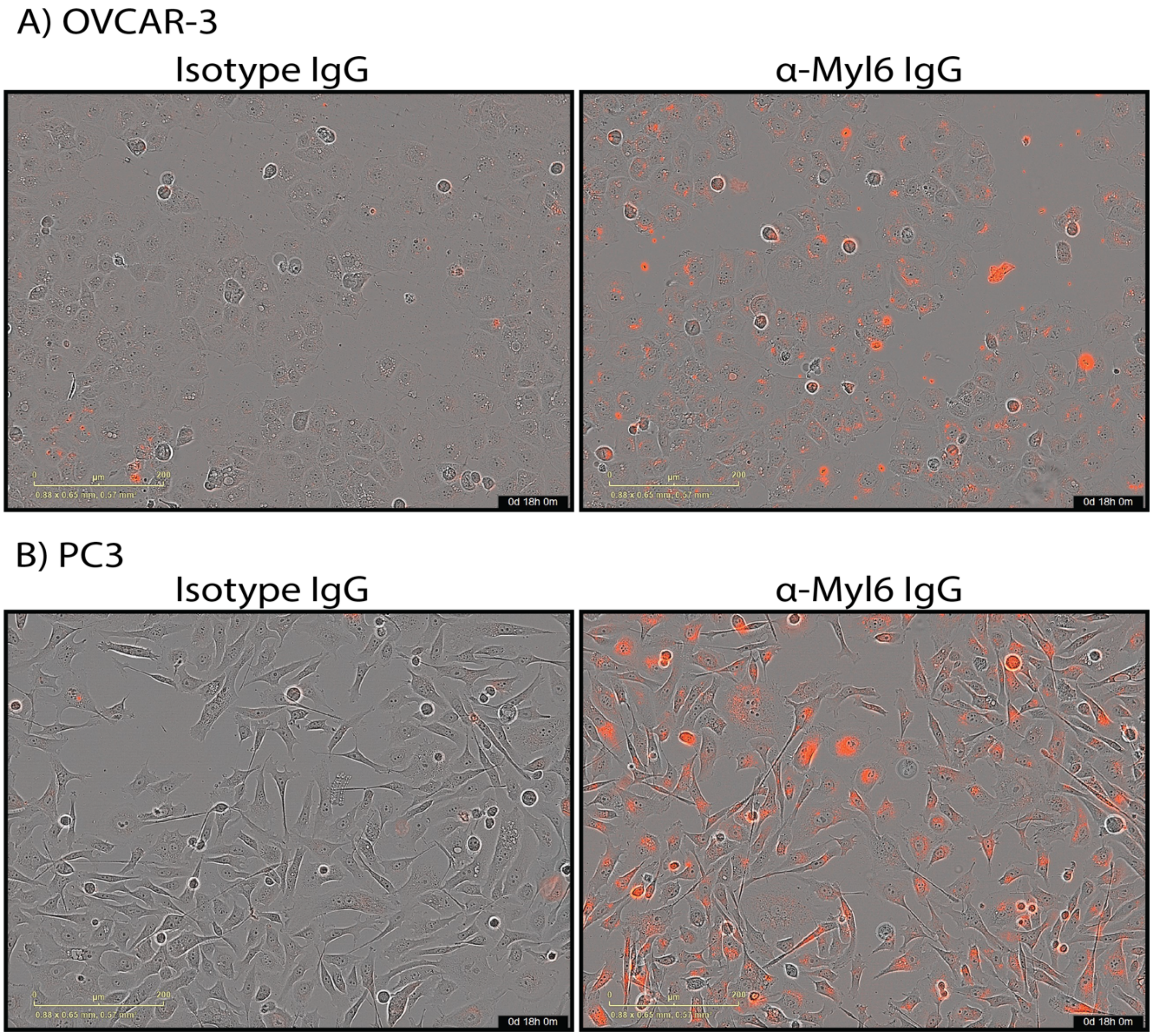


**Fig S10. Internalization of Myl6 in cancer cells monitored by real-time live-cell imaging.** OVCAR-3 and PC3 cells were subjected to real-time monitoring of anti-Myl6 IgG internalization using the Incucyte S3 live-cell analysis system. (**A**) The representative image of OVCAR-3 cells taken at 18 hours post-treatment with Isotype IgG (left) or anti-Myl6 IgG (right). No detectable red fluorescence is present in Isotype IgG, indicating the absence of IgG uptake, while substantial accumulation of intracellular red fluorescence is observed, reflecting robust internalization of anti-Myl6 IgG. (**B**) The representative image of PC3 cells taken at 18 hours post-treatment with Isotype IgG (left) or anti-Myl6 IgG (right). No detectable red fluorescence is present in Isotype IgG, indicating the absence of IgG uptake, while substantial accumulation of intracellular red fluorescence is observed, reflecting robust internalization of anti-Myl6 IgG.
